# Chronic Viral Reactivation and Associated Host Immune Response and Clinical Outcomes in Acute COVID-19 and Post-Acute Sequelae of COVID-19

**DOI:** 10.1101/2024.11.14.622799

**Authors:** Cole Maguire, Jing Chen, Nadine Rouphael, Harry Pickering, Hoang Van Phan, Abigail Glascock, Victoria Chu, Ravi Dandekar, David Corry, Farrah Kheradmand, Lindsey R. Baden, Rafick Selaky, Grace A. McComsey, Elias K. Haddad, Charles B. Cairns, Bali Pulendran, Ana Fernandez- Sesma, Viviana Simon, Jordan P. Metcalf, Nelson I Agudelo Higuita, William B. Messer, Mark M. David, Kari C. Nadeau, Monica Kraft, Chris Bime, Joanna Schaenman, David Erle, Carolyn S. Calfee, Mark A. Atkinson, Scott C. Brackenridge, Lauren I. R. Ehrlich, Ruth R. Montgomery, Albert C. Shaw, Catherine L. Hough, Linda N Geng, David A. Hafler, Alison D. Augustine, Patrice M. Becker, Bjoern Peters, Al Ozonoff, Seunghee Hee Kim-Schulze, Florian Krammer, Steve Bosinger, Walter Eckalbar, Matthew C. Altman, Michael Wilson, Leying Guan, Steven H. Kleinstein, IMPACC Network, Kinga K. Smolen, Elaine F. Reed, Ofer Levy, Holden Maecker, Peter Hunt, Hanno Steen, Joann Diray-Arce, Charles R. Langelier, Esther Melamed

## Abstract

Chronic viral infections are ubiquitous in humans, with individuals harboring multiple latent viruses that can reactivate during acute illnesses. Recent studies have suggested that SARS-CoV-2 infection can lead to reactivation of latent viruses such as Epstein-Barr Virus (EBV) and cytomegalovirus (CMV), yet, the extent and impact of viral reactivation in COVID-19 and its effect on the host immune system remain incompletely understood.

Here we present a comprehensive multi-omic analysis of viral reactivation of all known chronically infecting viruses in 1,154 hospitalized COVID-19 patients, from the Immunophenotyping Assessment in a COVID-19 Cohort (IMPACC) study, who were followed prospectively for twelve months. We reveal significant reactivation of *Herpesviridae*, *Enteroviridae*, and *Anelloviridae* families during acute stage of COVID-19 (0-40 days post-hospitalization), each exhibiting distinct temporal dynamics. We also show that viral reactivation correlated with COVID-19 severity, demographic characteristics, and clinical outcomes, including mortality. Integration of cytokine profiling, cellular immunophenotyping, metabolomics, transcriptomics, and proteomics demonstrated virus-specific host responses, including elevated pro-inflammatory cytokines (e.g. IL-6, CXCL10, and TNF), increased activated CD4+ and CD8+ T-cells, and upregulation of cellular replication genes, independent of COVID-19 severity and SARS-CoV-2 viral load. Notably, persistent *Anelloviridae* reactivation during convalescence (≥3 months post-hospitalization) was associated with Post-Acute Sequelae of COVID-19 (PASC) symptoms, particularly physical function and fatigue.

Our findings highlight a remarkable prevalence and potential impact of chronic viral reactivation on host responses and clinical outcomes during acute COVID-19 and long term PASC sequelae. Our data provide novel immune, transcriptomic, and metabolomic biomarkers of viral reactivation that may inform novel approaches to prognosticate, prevent, or treat acute COVID- 19 and PASC.

## Introduction

Viruses employ a variety of strategies to enhance their persistence and dissemination, including establishing chronic infection^1–3^. This strategy is exemplified by several human-infecting viruses, particularly those belonging to the *Herpesviridae* and *Anelloviridae* families, with these viruses establishing lifelong infections in a significant portion of the human population^4–6^. Although primary infection typically remains asymptomatic in immunocompetent individuals, some chronic viral infections contribute to the development of autoimmune disorders and cancers, among other adverse health outcomes^2,7–13^. These viruses typically remain dormant, but can reactivate during periods of stress, sleep deprivation, surgery, hormonal imbalances, or in the setting of critical illness^14–17^. The full range of immunological consequences stemming from these chronic viral reactivations remains largely unknown.

Since its emergence in 2019, Severe Acute Respiratory Syndrome Coronavirus 2 (SARS-CoV- 2) has resulted in over 774 million cases of coronavirus disease 2019 (COVID-19) and 7 million deaths^18,19^. Due to the physiological stress introduced by SARS-CoV-2 infection, underlying chronic viral infections may reactivate and potentially contribute to the immunological consequences of COVID-19. For example, reactivation of *Herpesviridae*, including EBV (Epstein-Barr Virus/Human Herpesvirus 4 (HHV4)), CMV (Cytomegalovirus/Human Herpesvirus 5 (HHV5)), Human Herpesvirus 6 (HHV6), and Human Herpesvirus 8 (HHV8), is associated with worse acute clinical outcomes in patients with COVID-19^20–24^. Reactivation of CMV and EBV, in particular, has been linked to more severe outcomes, including increased mortality^20,25^.

Additionally, patients with “Long COVID”, also known as Post-Acute Sequelae of COVID-19 (PASC), develop elevated EBV antibody titers, raising the possibility that reactivation of these viruses may contribute to PASC^22,26^.

Many of the foundational COVID-19 viral reactivation studies have been limited by small sample sizes, have focused on only a subset of *Herpesviridae*, or have relied solely on evaluating antibody responses to assess viral reactivation^20,22–24,26–30^, as opposed to measuring transcripts of actively replicating virus. As such, important gaps remain in our understanding of the dynamics and biology of chronic viral reactivation during acute COVID- 19, and their role in PASC.

To address the knowledge gap in viral reactivation in COVID-19, we leveraged samples and data from a longitudinal prospective observational study of 1,154 patients hospitalized for COVID-19 enrolled in the Immunophenotyping Assessment in a COVID-19 Cohort (IMPACC), with patients evaluated during acute hospitalization and for 12 months post hospital discharge. We carried out longitudinal, multi-omic analyses of nasal swabs, peripheral blood mononuclear cells (PBMCs), and endotracheal aspirates and found significant reactivation of chronic viruses, particularly from the *Herpesviridae* and *Anelloviridae* families, associated with acute COVID-19 severity. By integrating host and viral transcriptomics, cytokine profiling, cellular immunophenotyping, metabolomics, and proteomics, we observed distinct viral reactivation dynamics, and striking associations between viral reactivation, clinical outcomes, immunologic features, and patient demographics, both during acute COVID-19 and PASC. Our results provide novel insights into the endogenous virological landscape of COVID-19 patients, highlighting the complex interplay between SARS-CoV-2 infection, latent viral reactivation, host immune responses, and clinical outcomes.

## Results

### IMPACC Cohort

The IMPACC consortium enrolled 1,154 patients hospitalized for COVID-19 across 20 US hospitals between May 2020 and March 2021 (Figure 1A). All participants were COVID-19 vaccine-naive at the time of enrollment. To assess COVID-19 severity, participants were assigned to one of five trajectory groups (TG) using latent class mixed modeling of respiratory status over the first 28 days^31^. Groups were classified as mild (TG1), moderate (TG2), severe (TG3), critical (TG4), or fatal within 28 days (TG5). From each participant, bulk RNA sequencing was performed on PBMCs, nasal swabs, and for mechanically ventilated patients, endotracheal aspirates (EA), at up to ten visits during one-year post-hospital admission (Figure 1B). In addition, we assessed whole blood immune cell populations by mass cytometry by time of flight (CyTOF), serum anti-EBV and anti-CMV antibody titers, serum cytokine levels by proximity extension assay (PEA), and the plasma proteome and metabolome by mass spectrometry at participant visits.

**Figure 1.**
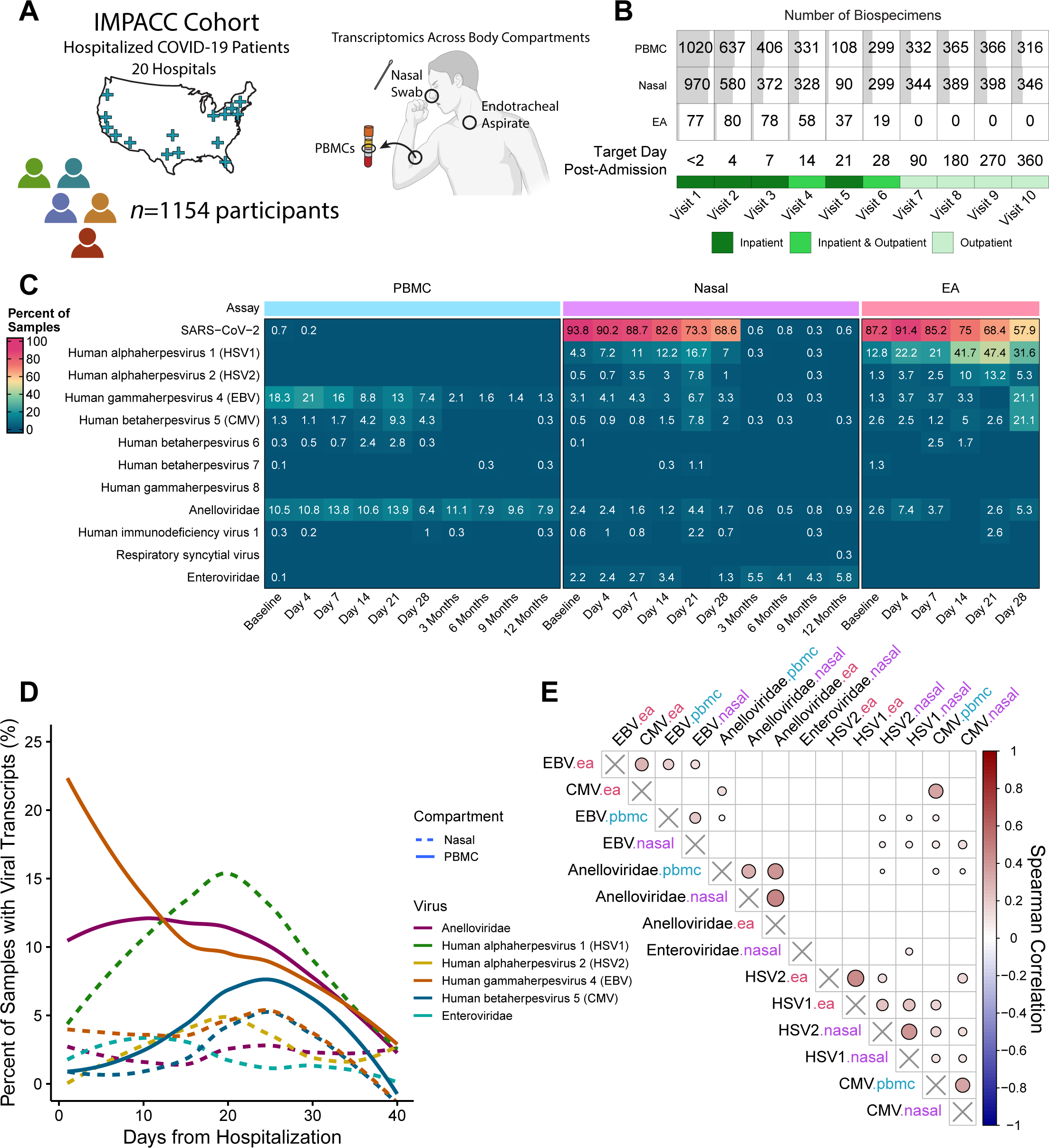
The Transcriptionally Active Human Virome of the Blood, Upper Airway and Lungs in Patients Hospitalized for COVID-19. **A)** A total of 1154 participants were recruited across 20 sites for the IMPACC study, with PBMC, nasal, and EA samples analyzed with RNA-sequencing. **B)** The number of participants with the respective transcriptomic at each timepoint. The background gray shading indicates the percentage of total participants, with samples analyzed at that timepoint for each transcriptomic assay. **C)** Heatmap of the percent of samples that had detected reads for various latent viruses for each transcriptomic assay at the different timepoints. **D)** Smoothed curves demonstrating the proportion of total samples that were positive for the five most common viruses in the nasal and PBMC transcriptomics. Curves were calculated by the proportion of samples positive for each day ± two days (a rolling window approach), followed by a local polynomial regression fitting. **E)** Spearman correlation of viral reads per million across transcriptomic assays, with viruses hierarchically clustered. Correlation in (E) only visualized if adj.p <= 0.05. Source data are provided as a Source Data file.

### RNA-sequencing Identifies Transcripts from the Human Virome

From RNA-seq data, we identified viral RNA transcripts in nasal, EA and PBMC samples and found a diverse number of human infecting viruses beyond SARS-CoV-2 including EBV, CMV, HHV6, HSV1, HSV2, and several *Anelloviridae* and *Enteroviridae* species (Figure 1C).

Unsurprisingly, SARS-CoV-2 was the most prevalent virus identified, and was primarily found in NS and EA samples, with detection in PBMCs only in 10 participants near time of admission (Figure 1C). We confirmed that SARS-CoV-2 abundance measured by RNA-seq reads per million (rpM) highly correlated with RT-qPCR cycle threshold (Supplemental Figure S1A-B).

Among *Herpesviridae*, HSV1/2, EBV, and CMV transcripts were commonly detected across compartments during acute COVID-19 (defined as the first 40 days after hospital admission), with a notable lack of detection during the convalescent period (>3 months post admission) (Figure 1C). In addition, we also detected a diverse number of *Anelloviridae* and *Enteroviridae* species (Figure S1C-D), with their collective viral load at the family taxonomic level used for analyses.

Interestingly, we found that each viral species displayed unique temporal dynamics of reactivation relative to hospital admission (Figure 1D). For example, EBV reactivated early in the disease course, with ∼20% of participants having detectable transcripts at the time of admission, followed by a gradual decline in detection over time. Unlike EBV, the frequency of *Anelloviridae* transcripts remained constant up to day 20 post-admission, followed by a slow decline. In contrast, HSV1/2 and CMV reactivated later in disease, with HSV1 detected in up to 40% of EA samples and CMV in ∼8% of PBMC samples 19-24 days post admission (Figure 1D & S1D-E).

The detection of each virus varied across compartments, with EBV transcripts more common in PBMCs and HSV1/2 transcripts notably more common in NS and EA samples. Evaluation of viral rpM across nasal, EA and PBMC compartments demonstrated that transcript abundance for individual viruses was often correlated across the three compartments (Figure 1E, Supplemental Figure S1F).

### Activation of the Human Virome is Associated with COVID-19 Clinical Outcomes

Next, we evaluated how detection of viral transcripts in the first 40 days post hospital admission is associated with COVID-19 severity, using the previously published IMPACC trajectory groups (TG)^31^ (Figure 2A, and S2A, and Supplementary Data 1). Cumulative linked modeling of the TGs demonstrated significant associations between COVID-19 severity and the detection of *Herpesviridae* and *Anelloviridae* transcripts (Supplementary Data 1). More specifically, we found associations between severity and the detection of transcripts from *Anelloviridae* (PBMC adj.p = 2.37E-05), CMV (nasal adj.p = 3.16E-04, PBMC adj.p = 9.91E-04), EBV (nasal adj.p = 7.33E-06, PBMC adj.p =8.06E-10), HSV1 (nasal adj.p =1.05E-05), and HSV2 (nasal adj.p = 6.61E-04 transcripts. When further limiting to severely ill TG4 patients (still hospitalized after 28 days), we observed that patients with CMV transcripts in any compartment were more likely to die within one year (nasal adj.p = 4.79E-03, EA adj.p = 6.66E-03, PBMC adj.p = 6.66E-03). This was also the case for patients with detectably expressed EBV transcripts in the upper respiratory tract (adj.p = 0.0067), HSV1 (adj.p = 0.0067), or HSV2 (adj.p = 0.0067) (Figure 2A). Of note, there was no difference in prevalence of chronic viruses between TG4 and TG5.

**Figure 2.**
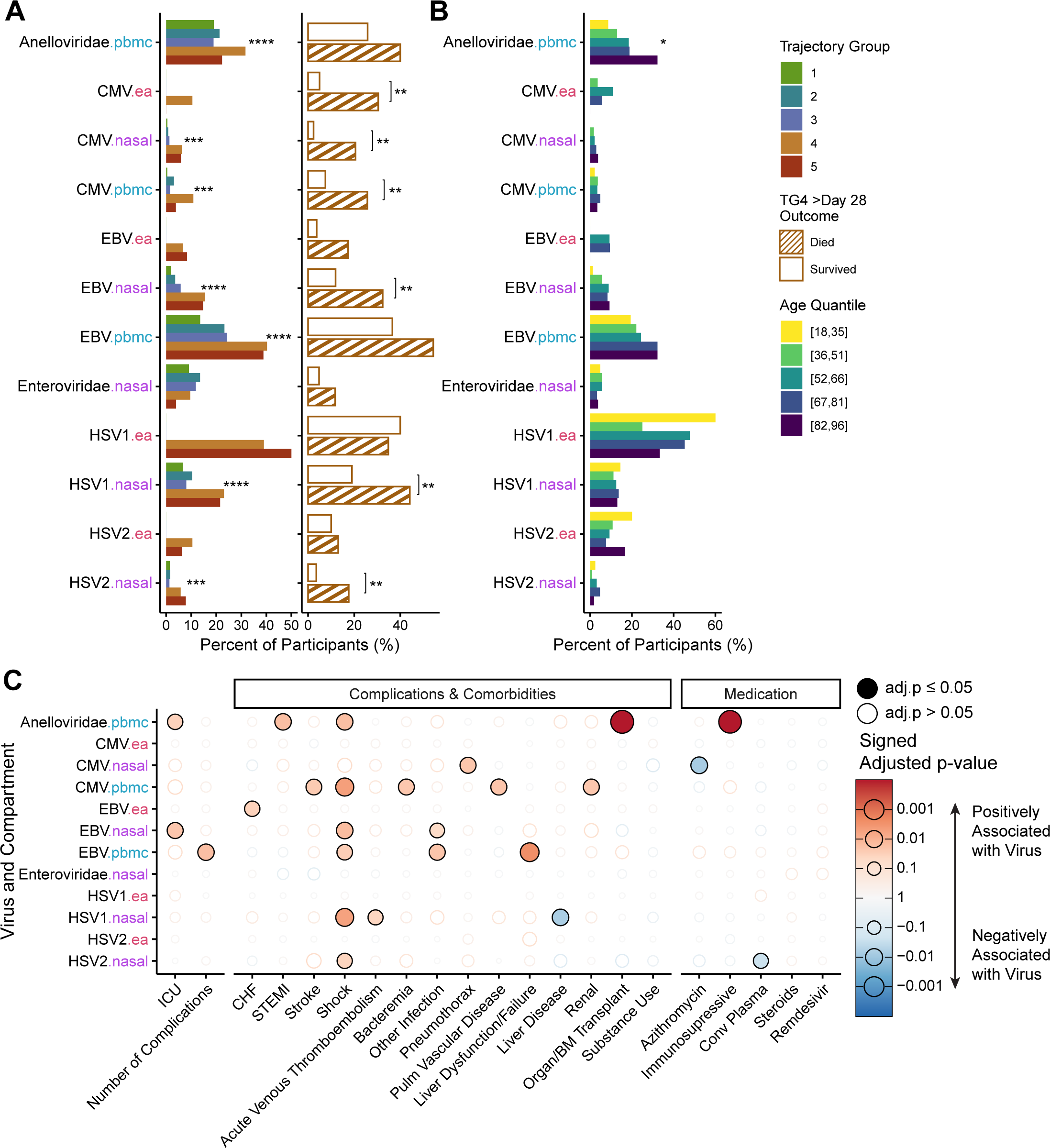
Clinical Outcomes Associated with Activation of the Human Virome in Severe COVID-19. **A)** Percent of participants in the cohort who had detectable viral reads in at least one sample within 40 days of hospital admission for the respective virus and tissue split by each trajectory group (on the left), a measure of COVID-19 severity, and participants in TG4 split by their long-term mortality outcome (on the right). **B)** Percent of participants in the cohort who had detectable viral reads in at least one sample within 40 days of hospital admission for the respective virus and tissue across age groups (split by quantiles). In (A) and (B) adjusted p-values of <0.05, <0.01, <0.001, and <0.0001 are represented by *, **, ***, and **** respectively. **C)** Results from generalized linear mixed modeling of various complications, comorbidities, and medication usage evaluating for association with viral transcripts detected within 40 days of hospitalization while controlling for sex, age quantile, and TG. Directionality of the association indicated by color, with a positive association increasingly red and a negative association increasingly blue. (Filled dots represent adj.p < 0.05, p-values adjusted using Benjamini-Hochberg procedure). Source data are provided as a Source Data file.

We then evaluated whether the detection of chronically infecting viral transcripts varied with age, while adjusting for COVID-19 severity (TGs), and found a significant positive association with *Anelloviridae* transcripts in the PBMCs with increasing age (Figure 2B, adj.p = 0.044, Supplementary Data 1). When assessing associations between race or ethnicity, we found that Hispanic ethnicity was significantly associated with detection of both CMV (adj.p = 0.03) and EBV transcripts (adj.p = 0.03, Figure S2B, Supplementary Data 1). However, we found no significant associations between viral transcripts and biological sex, or treatment with either remdesivir or steroids (Figure S2C-E).

To further extend our analysis, we also evaluated the association of viral transcripts with comorbidities, medication usage, and complications (Figure 2C, Supplementary Data 1). *Anelloviridae* transcripts in PBMCs were significantly associated with a history of solid organ transplantation and immunosuppression, as well as shock and ST-elevation myocardial infarction (STEMI). CMV in the nasal compartment was linked to pneumothorax, whereas CMV in the PBMCs was associated with bacteremia, pulmonary vascular disease, renal complications, shock, and stroke. Interestingly, CMV in the nasal compartment was also inversely associated with azithromycin use. EBV transcripts in the nasal compartment were primarily associated with ICU-level care and shock, while detection of EBV transcripts in PBMCs correlated with liver failure, concurrent infections, shock, and the overall number of complications. Also, detection of EBV transcripts in the nasal compartment and the PBMCs was associated with a secondary infection besides SARS-CoV-2. Detection of HSV1 in the nasal compartment had a significant association with acute venous thromboembolism and shock and was inversely associated with liver disease. Similarly, HSV2 reads in the nasal compartment were associated with shock and inversely associated with convalescent plasma.

### Viral Reactivation Results in Elevated Antibody Titers and Changes in Immune Cell Frequencies

We next leveraged our multi-omic data to further validate chronic viral reactivation in COVID-19 and characterize the associated responses of the host’s immune system, incorporating serum EBV and CMV antibody levels, immune cell frequencies, serum cytokines, plasma metabolomics, and host transcriptomics.

First, we observed that patients with detectable EBV transcripts in PBMCs had persistently elevated EBV IgG and IgA antibody titers (Figure 3A, Supplementary Data 2). Similarly, patients with CMV transcripts had significantly higher CMV seropositivity rates at baseline (Figure 3B, Supplementary Data 2). Furthermore, using unbiased mass spectrometry proteomics^32,33^, we assessed circulating HSV1 proteins in participant plasma. These proteins were more common in participants from whom HSV1 transcripts were identified in nasal swabs (Figure 3C, p=0.02, Supplementary Data 2), and significantly more common in TG4 patients with severe COVID-19 (Figure S3B, p=0.0003, Supplementary Data 2).

**Figure 3.**
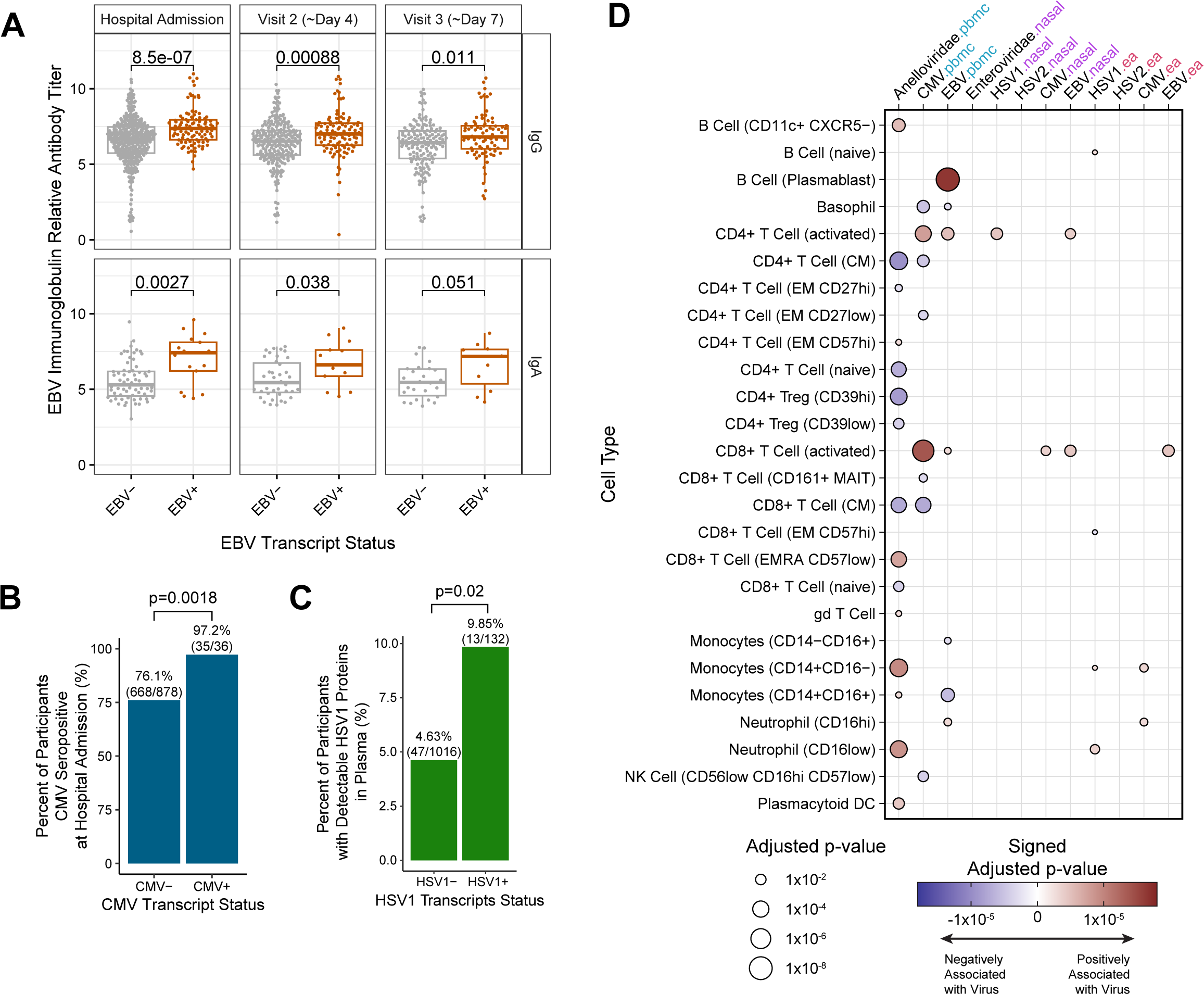
Antibody Response, Proteomics, and Circulating Cellular Immunophenotyping Validates Chronic Viral Reactivation in COVID-19. **A)** Relative EBV GP350 IgG Antibody Titers in participants with detected EBV transcripts (EBV+) vs participants with no detected transcripts (EBV-). Adjusted p-values calculated using a Wilcoxon rank-sum test with Benjamini-Hochberg corrections. **B)** Percent of patients seropositive for CMV at admission in participants with no CMV transcripts in the acute period (CMV-) vs participants with detectable transcripts (CMV+). **C)** Percent of participants with detectable HSV1 proteins in the plasma for participants with HSV1 detectable in the nasal swabs (HSV1+) vs participants with no detectable transcripts (HSV-). P-values in (B) and (C) calculated using a chi-square test of independence. **D)** Results of linear mixed effect modeling identifying whole blood cell frequencies significantly associated with detection of viral reads for different viruses while controlling for TG, sex, and age. Directionality of the association indicated by color, with a positive association increasingly red and a negative association increasingly blue. (Dot only shown if adj.p <= 0.05.). Source data are provided as a Source Data file.

Using mass cytometry by time of flight (CyTOF), we next evaluated relationships between viral transcript detection and blood immune cell frequencies. We observed significant associations between EBV transcription in PBMCs and increased proportions of B-cell plasmablasts, the primary host cells of the virus (Figure 3D, Supplementary Data 2). Furthermore, we observed that detection of CMV transcripts was associated with a significant reduction in the frequency of CD4 and CD8 central memory T cells, CD27low effector memory CD4 T cells, and CD56low CD16hi CD57low NK cells. Interestingly, detectable expression of both EBV and CMV was associated with a higher frequency of activated CD4+ and CD8+ T cells.

To assess whether changes in circulating cell frequencies may explain the association between *Herpesviridae* and *Anelloviridae* transcripts and COVID-19 severity we repeated our analysis of viral transcripts across the TGs while controlling for immune cell frequencies, which vary with disease severity^34^. This analysis demonstrated that even when controlling for changes in these underlying cell frequencies, viral transcripts were still significantly associated with increasing COVID-19 severity (Figure S3A, Supplementary Data 2).

### Reactivation of the Human Virome Correlates with Changes in Inflammatory Cytokines and Chemokines

Next, we asked whether detection of *Herpesviridae* and *Anelloviridae* transcripts was associated with changes in inflammatory protein levels. Using generalized additive mixed modeling (gamm), we compared the longitudinal dynamics of cytokines in patients with or without evidence of viral reactivation, while controlling for COVID-19 severity (TGs), sex, and age (Figure 4A, Supplementary Data 3).

**Figure 4.**
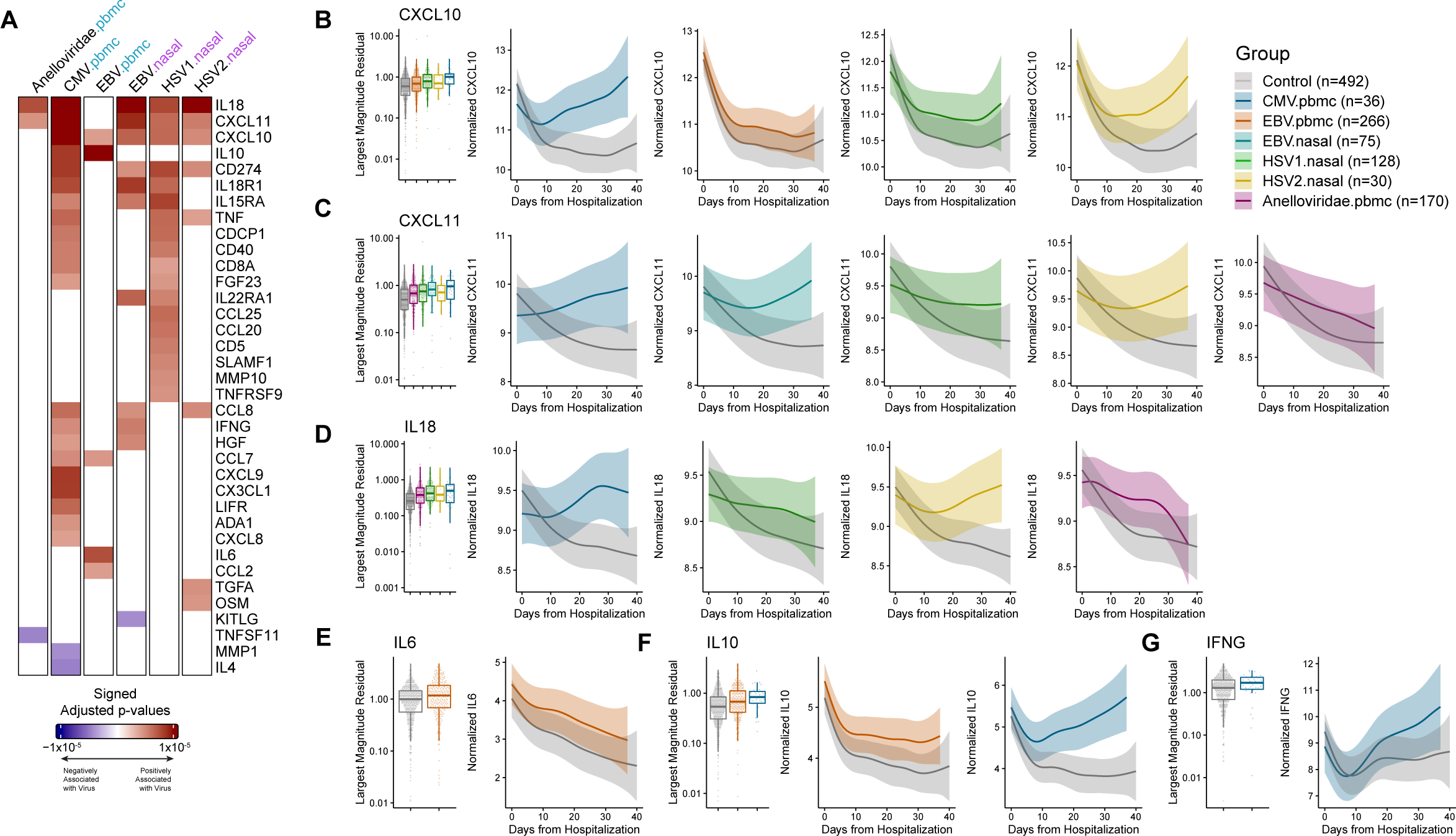
Activation of the Human Virome Correlates with Changes in Inflammatory Cytokine Expression. **A)** Summary heatmap of adjusted p-values colored by direction of significance for associations of cytokines with specific viruses in the upper respiratory tract and blood. Adj.p shown if adj.p<0.01 for either the main effect or time interaction term in the model. Boxplot of largest gamm residuals for each participant by virus and longitudinal 95% confidence interval for **B)** CXCL10, **C)** CXCL11, **D)** IL18, **E)** IL6, **F)** IL10, and **G**) IFNG. Source data are provided as a Source Data file.

Interestingly, we found reactivation of different chronic viruses associated with several unique cytokine, chemokine, and inflammatory soluble protein signatures (Figure 4A and S4). However, there was also a set of shared cytokines including CXCL10 (Figure 4B), CXCL11 (Figure 4C), and IL18 (Figure 4D) that were correlated with several chronic viruses.

EBV in the PBMC was associated with elevation of several key cytokines including IL6 (Figure 4E), CCL7, CCL2, IL10 (Figure 4F), and CXCL10 all of which have been associated with COVID-19 severity^35–37^ (Figure 4A and S4). Interestingly, from the cytokines associated with EBV in PBMC, only CXCL10 was also associated with EBV in the nasal transcriptomics. In addition, EBV in the nasal compartment was also correlated with increases in IL18, CXCL11, CXCL10, CD274, IL18R1, IL15RA, IL22RA1, CCL8, IFNG, HGF, and a decrease in KITLG.

Thus, these data suggest that the host immune response towards chronic viruses may differ between different compartments, with EBV in the nasal compartment possibly reflecting a more severe state of EBV viral reactivation.

Similar to EBV, the other detected herpetic viruses (HSV1 and HSV2 in the nasal and CMV in the PBMC transcriptomics) associated with pro-inflammatory serum cytokines, with many in common between the viruses. HSV1, HSV2, and CMV were all associated with increases in IL18, CXCL11, CXCL10, CD274, and TNF, with HSV1 and CMV also correlated with elevations in IL18R1, IL15RA, CDCP1, CD40, CD8A, FGF23. Furthermore, HSV2 was associated with increases in CCL8, TGFA, and OSM, HSV1 with elevations of IL22RA1, CCL25, CCL20, CD5, SLAMF1, MMP10, and TNFRSF9, and CMV with increases in CCL8, IFNG (Figure 4G), HGF, CCL7, CXCL9, CX3CL1, LIFR, ADA1, CXCL8, and decreases in MMP1 and IL4. Finally, *Anelloviridae* were only associated with an increase in CXCL11 and IL18, and a decrease in TNFSF11. We also observed compartment-specific differences in the relationship between cytokine expression and viral replication. For instance, of the cytokines associated with EBV transcripts in PBMCs, only CXCL10 was also associated with EBV in the upper airway.

### Reactivation of the Human Virome Correlates with Changes in the Metabolome

Using the same longitudinal gamm analysis, we evaluated the association of chronic viral reactivation with plasma metabolites as measured by liquid chromatography-mass spectrometry (Supplementary Data 4). As has been observed with other viral infections^38^, reactivation was associated with changes in metabolites belonging to amino acid and lipid metabolism^38^ (Figure 5A and 5B). Of the viruses with detectably expressed transcripts, CMV was associated the greatest number of changes in the metabolome, and in particular with higher levels of urea and TMAP, metabolites previously linked to kidney injury (Figure 5A, 5C and S5B)^39,40^. Additionally, detection of CMV and *Anelloviridae* transcripts were both associated with increases in several long chain fatty acids such as erucate, arachidate, and docosadienoate (Figure 5D and S5C-D) and regulators of nitric oxide synthesis, dimethylarginine (SDMA + ADMA)^41–44^ (Figure S5E). We also identified a shared metabolomic signature common across multiple viruses (Figure S5A).

**Figure 5.**
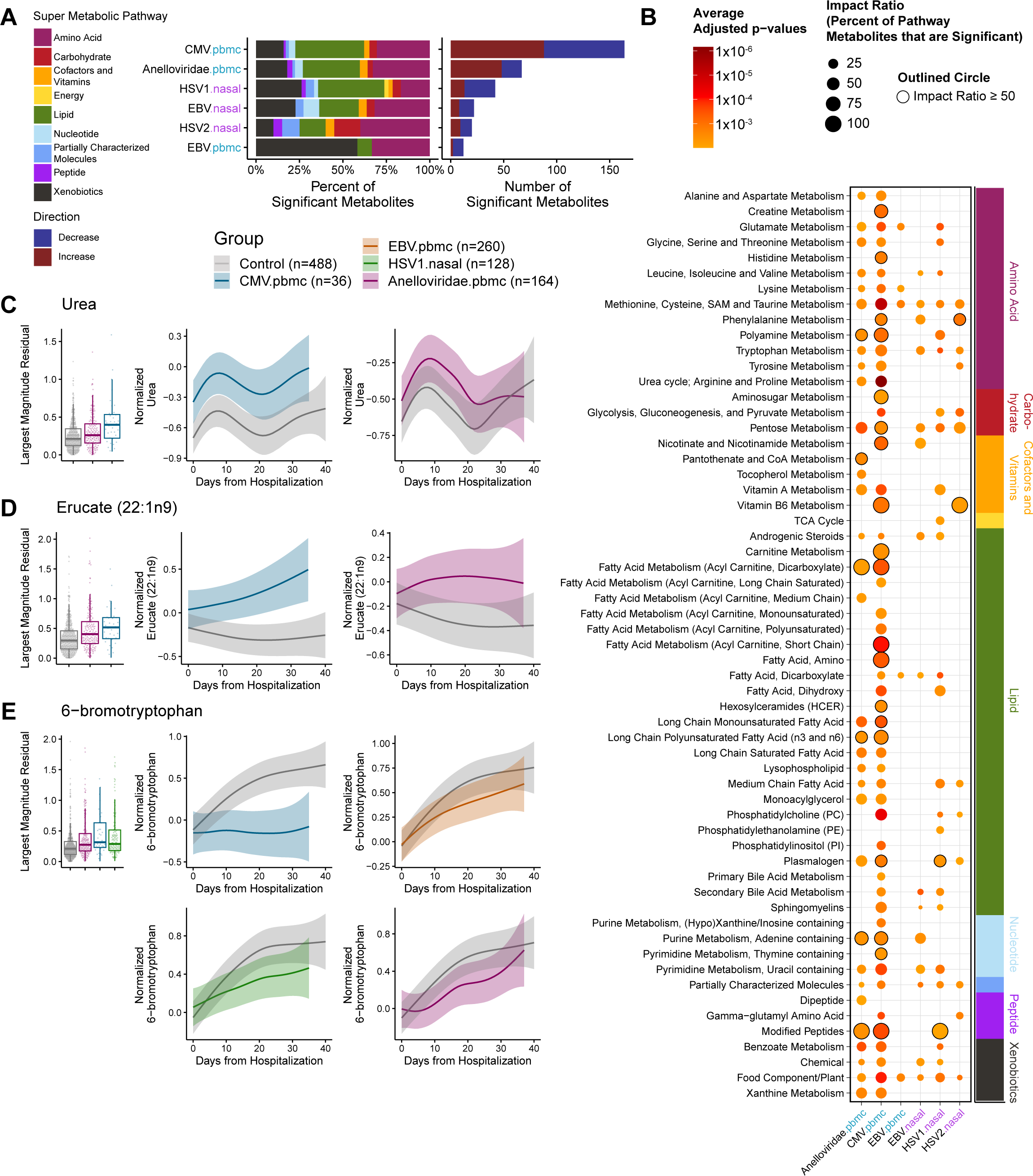
Viral reactivation associated with shifts in the plasma metabolome. **A)** Number of significant metabolites for each chronically infecting virus across tissues and the percentage of significant metabolites that map to each major branch of metabolism. **B)** Dot plot of metabolic sub-pathways containing significantly different metabolites for the different viruses, with dot size indicating the impact ratio (the percent of detected metabolites from that pathway that were significantly different) and the color indicating the average adjusted p-value of significant metabolites in that pathway. Largest magnitude gamm residual for each participant and longitudinal 95% confidence interval for **C)** urea, **D)** erucate, and **E)** 6-bromotryptophan. Source data are provided as a Source Data file.

For example, both S-methylcysteine sulfoxide and 6-bromotryptophan were reduced with reactivation of *Anelloviridae*, HSV1, HSV2, CMV, and EBV (Figure 5E and S5F).

### Reactivation of Chronic Viruses is Associated with Changes in the Host Transcriptome

To identify a signature for each chronic virus independent of COVID-19 severity and participant demographics, we evaluated nasal and PBMC transcriptomic data for differentially expressed host genes while controlling for COVID-19 severity, SARS-CoV-2 nasal viral load, sex, age, and days from hospital admission (Supplementary Data 5).

In the PBMC transcriptomics, detection of viral transcripts in both the PBMC or nasal compartments were associated with changes in host PBMC gene expression (Figure 6A and S6, Supplementary Data 5). Hypergeometric enrichment analysis demonstrated that *Anelloviridae* in the PBMCs, CMV in the PBMCs, and HSV1/2 in the nasal compartment were all associated with downregulation of a diverse number of pathways pertaining to RNA processing and protein translation. Similarly, EBV, CMV, HSV1/2 were associated with an upregulation of pathways pertaining to cellular replication. In addition to these shared pathways, *Anelloviridae* in the PBMCs, CMV in the PBMCs, and HSV1/2, and CMV in the nasal compartment were also strongly associated with signatures of neutrophil degranulation. Finally, EBV in the PBMCs was uniquely associated with platelet activation and signaling.

**Figure 6.**
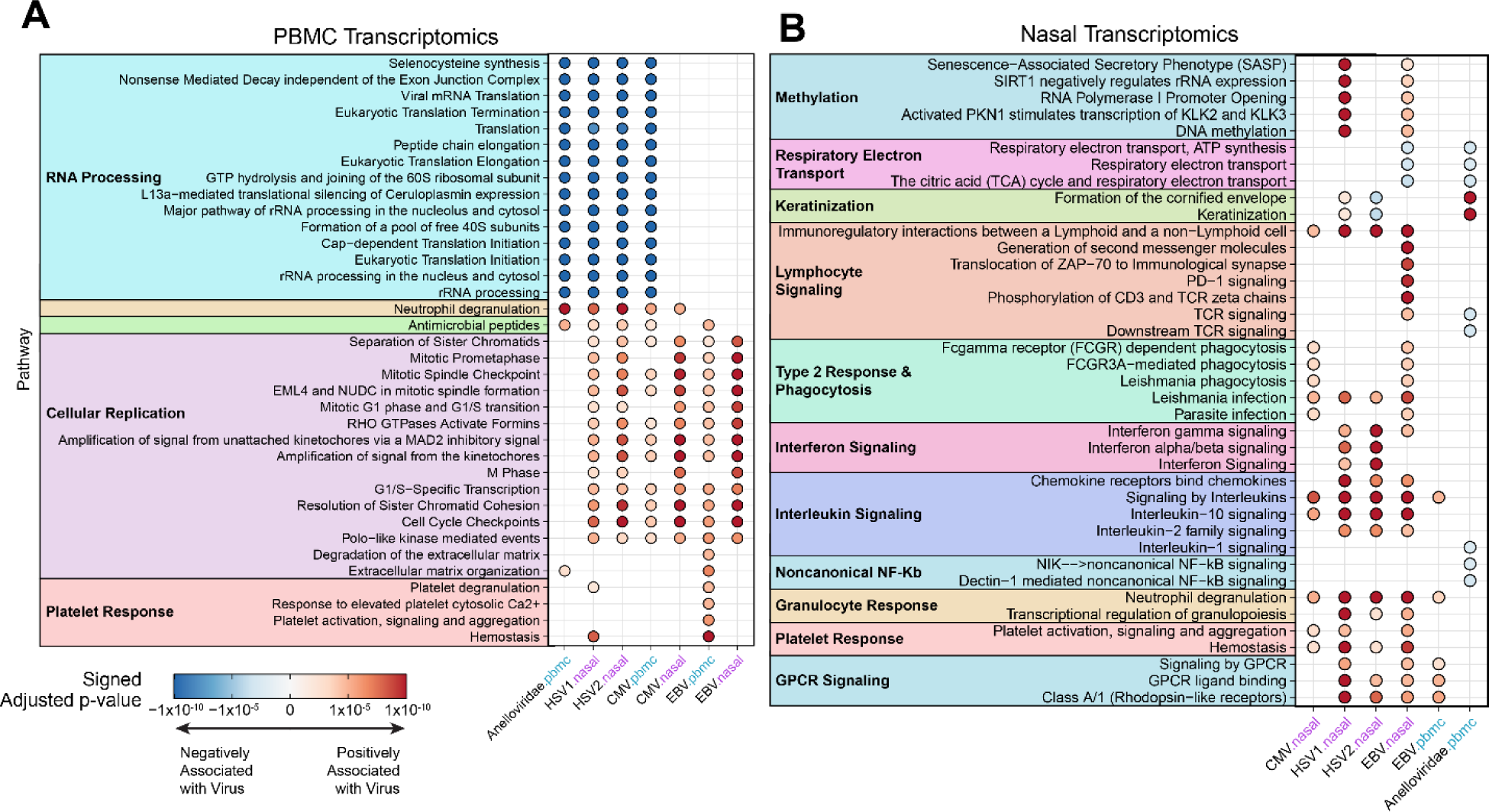
Chronic Viral Reactivation is Associated with Alterations in Host Gene Expression. **A)** Top enriched pathways in the PBMC transcriptomics associated with reactivation of the identified latently infecting viruses. **B)** Top enriched pathways in the nasal transcriptomics associated with reactivation of the identified latently infecting viruses. Results for (A) and (B) calculated using hypergeometric enrichment of pathways from Reactome, separately on the positive and negative differentially expressed genes. Dot only shown for a pathway if adjusted p-value < 0.01. Source data are provided as a Source Data file.

When evaluating the nasal transcriptomics, viruses that reactivated in the nasal compartment (EBV, CMV, and HSV1/2) had the strongest associations with changes in upper airway gene expression (Figure 6B and S6, Supplementary Data 5), with inflammatory signaling pathways significantly upregulated in patients with reactivation of these viruses. For example, CMV, EBV, and HSV1/2 transcripts were all associated with upregulation of interleukin signaling, lymphocyte immunoregulatory interactions, and neutrophil degranulation. Of note, interleukin signaling included Interleukin-10 signaling, which was elevated in the serum of participants with CMV and EBV reactivation. Additionally, CMV and EBV replication in the upper airway was also associated with increases in pathways pertaining to a type 2 immune response and phagocytosis. Detection of HSV1 or EBV transcripts in the nasal compartment was associated with upregulation of methylation, and EBV was specifically associated with increases in pathways pertaining to T-cell activation. Finally, *Anelloviridae* in the PBMCs was also associated with changes in the nasal transcriptome, with the strongest associations pertaining to upregulation of genes involved in keratinization and downregulation of noncanonical NF-Kb signaling.

### Detection of Anelloviridae Transcripts Associates with Physical Disability and Fatigue in PASC patients

Given recent reports linking chronic viral reactivation to PASC^22,26,27^, we evaluated how viral reactivation correlated with our previously identified patient reported outcome (PRO) groups using convalescent survey data^45^. The three PASC groups consisted of physical (characterized by physical disability and fatigue), cognitive (characterized by cognitive impairment), and global deficits (characterized by both physical and cognitive deficits). These groups were compared to a fourth PRO group reporting minimal deficits (minimal). To probe the relationship between chronic viral reactivation and PASC PRO groups, we examined whether viral reactivation was more prevalent in specific PRO groups during either acute or convalescent stages of COVID-19.

Upon evaluating the relationship between viral reactivation during the *acute* stage of COVID-19, no significant relationship was found with the PRO groups (Figure 7A, Supplementary Data 6). However, due to the association of chronic viruses with mortality and the participant drop-out in the convalescent stage, the sample size was limited (Figure S7A, Supplementary Data 6).

**Figure 7.**
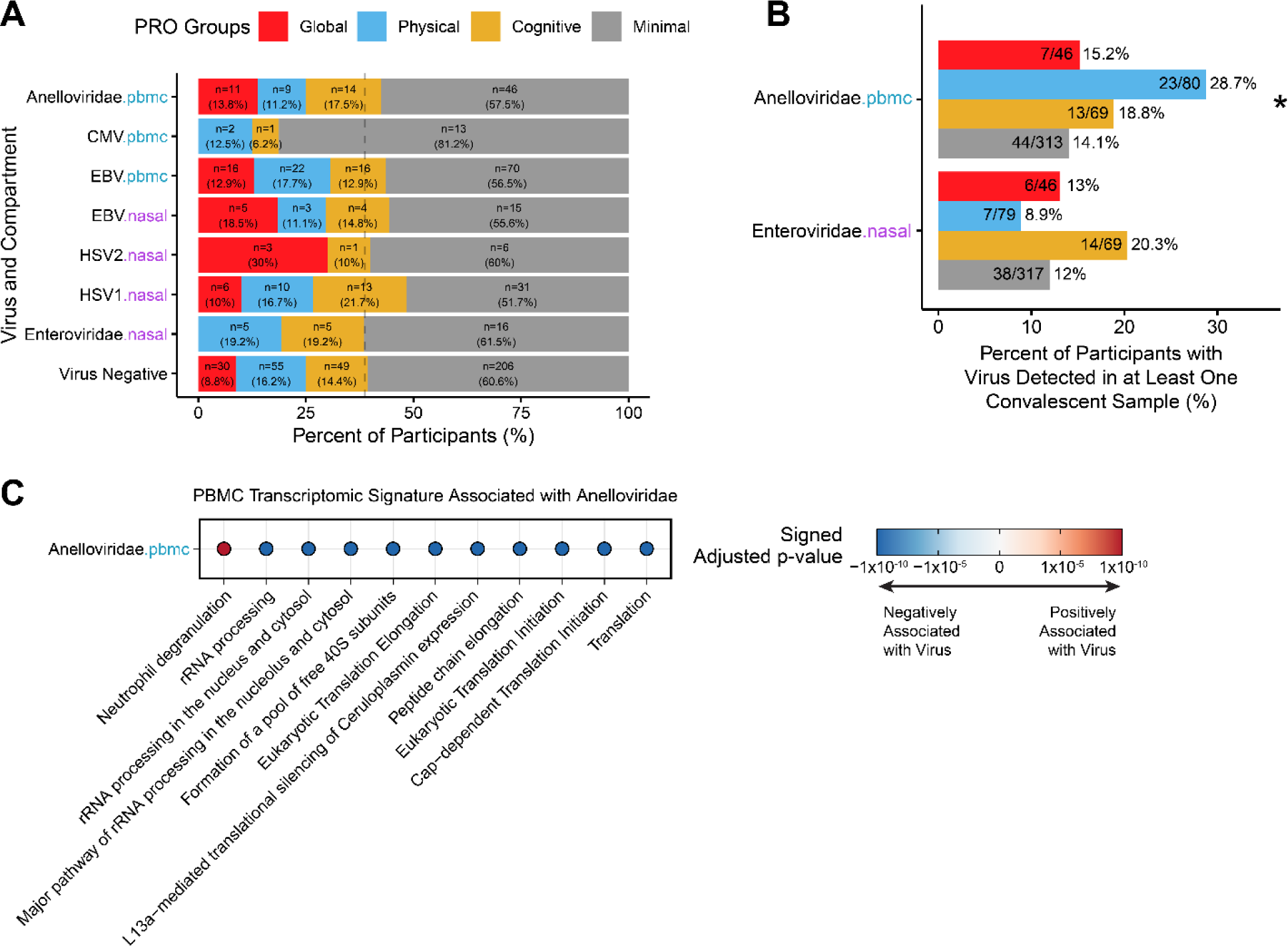
Associations between Virome Activation and PASC. **A)** Percent of participants with reactivation of chronically infecting viruses across specific compartments (nasal or PBMC) that fall into PASC associated patient reported outcome (PRO) groups. The virus negative group are participants who had no viral reactivation for any virus in the acute period, whose rate of total PASC is indicated by the gray line and serves as the baseline reference. **B)** Percent of the PRO groups which had viral transcripts detected for *Anelloviridae* in the PBMCs, or *Enteroviridae* RNA detected in the upper airway, in any of their convalescent samples. Significance calculated using chi-square test, * indicates adjusted p-value < 0.05. **C)** Top hypergeometric enriched pathways (determined via lowest adjusted p-values) from DEGs associated with *Anelloviridae* using convalescent samples reveals a similar signature of *Anelloviridae* during acute COVID-19 (shown in Figure 6A). Source data are provided as a Source Data file.

Despite the lack of statistical significance during the acute stage, the relationship between specific viruses and PASC trended in the direction of previous reports, with CMV reactivation associated with lower rates of PASC ^22^.

We then investigated the correlation between viral detection during the *convalescent* period (> 60 days post hospitalization) and PRO groups. Interestingly, among the examined viruses, *Anelloviridae* and *Enteroviridae* were the most frequently detected viruses in RNA-seq of convalescent samples (Figure 1C and 7B, Supplementary Data 6). Notably, we found that *Anelloviridae* transcripts were significantly more prevalent in participants from the Physical PRO group (*X^2^* = 9.95, adj.p = 0.038), which was characterized by high scores on the Patient-Reported Outcomes Measurement Information System (PROMIS) Physical Function survey.

This finding suggests that detection of *Anelloviridae* transcripts may serve as a potential biomarker for persistent physical disability in PASC patients. We further observed the same *Anelloviridae* gene expression signature during both the acute and convalescent periods (i.e. an upregulation of genes involved in neutrophil degranulation and downregulation of genes involved in RNA processing, Figure 6A and 7C).

## Discussion

Chronically infecting viruses are prevalent in the general population, with individuals estimated to carry an average of 8 to 12 chronic viral infections at any given time^2^. These viruses are generally innocuous and typically do not result in acute infectious symptoms, however, emerging evidence suggests that their reactivation is associated with and may contribute to multiple diseases, including autoimmune syndromes, malignancies, and chronic fatigue syndrome ^2,7–13,46^. Additionally, *Herpesviridae*, such as EBV and CMV, have been found to reactivate in severe infections and sepsis^16,47,48^ and have more recently been implicated in the pathophysiology of Long COVID/PASC^22,26,27^. Thus, there is an urgent need to better understand how chronically infecting viruses reactivate in COVID-19 and to develop preventative and therapeutic strategies to combat their reactivation.

Here, we conducted a prospective, multi-omic analysis of blood and respiratory samples from >1000 hospitalized COVID-19 patients, which revealed widespread reactivation of chronic viral infections, specifically from the *Herpesviridae* and *Anelloviridae* families. By integrating data from multiple platforms, including cellular and cytokine immunophenotyping, metabolomics, and transcriptomics, our findings expand on the complex interplay between SARS-CoV-2 infection and chronic viral reactivation, and provide a deeper understanding of the host immune response. Importantly, our findings demonstrate association of viral co-infections on the clinical course of COVID-19 and PASC.

Despite prior studies evaluating viral reactivation in acute COVID-19^20,21,25^, the exact timing of reactivation for different viruses has not been clearly established. Here, leveraging a large and longitudinally sampled cohort, we define timing and duration of viral reactivations. Specifically, we found that EBV transcripts in blood peaked early in acute disease following hospital admission, and then decreased over time. Similarly, detection of *Anelloviridae* transcripts from blood was most common early during hospitalization and began to decline several weeks later. In contrast, CMV, HSV1, and HSV2 displayed later reactivation and were found primarily in respiratory samples, peaking at approximately 22 days post hospitalization. Interestingly, patients who were EBV transcript positive, in either the nasal or PBMC compartments at hospital admission, had higher relative EBV IgG antibody titers, suggesting that EBV reactivation may happen prior to hospitalization for some patients. This may also be the case for HSV1, HSV2, and CMV as reactivation of these viruses may be present in tissues before they become detectable in PBMC or mucosal secretions. These results provide novel insights into the dynamics of the diverse virological landscape of COVID-19 and reveal that the time course of reactivation varies for different viruses.

Reactivation of *Herpesviridae* and *Anelloviridae* was associated with key clinical outcomes, including severe disease and death. CMV reactivation in all tested compartments (upper and lower respiratory tract and blood) was associated with increased mortality; while EBV reactivation in the nasal compartment and HSV1/2 reactivation in the respiratory compartment served as prognostic markers of mortality. Renal complications were more common with CMV reactivation, while venous thromboembolism was most associated with HSV1.

Immunocompromised patients were most likely to experience *Anelloviridae* reactivation, as previously reported^49–51^. Shock was associated with multiple viral reactivations across anatomical compartments including *Anelloviridae* in PBMCs, CMV and EBV in the upper respiratory tract and PBMCs, and HSV1/2 in the upper respiratory tract, while EBV emerged as the virus with the most significant association to the overall number of complications.

Interestingly, azithromycin, a macrolide antibiotic, known to have antiviral activity^52^ and that exhibits anti-inflammatory effects that could influence microbial infections^53,54^, was associated with a decrease in CMV prevalence in the nasal compartment. These findings collectively raise the possibility of a multifaceted role of viral reactivation in the exacerbation of acute COVID-19, underscoring the need for integrated viral surveillance across body compartments and integration of viral reactivation data into the management of acute COVID-19 patients.

Regarding demographic features associated with virome reactivation, our findings are in line with prior epidemiological studies demonstrating higher rates of reactivation for EBV and CMV in individuals of Hispanic/Latino ethnicity^55^. Detection of *Anelloviridae* transcripts was significantly associated with increasing age, aligning with previous findings of higher *Anelloviridae* viral loads in older adults^56^. This finding suggests that aside from the risk of more severe disease, age on its own may not be a critical factor in herpetic viral reactivations.

Interestingly, there was a unique signature of cytokines, chemokines, and other inflammatory proteins associated with reactivation for the various viruses. For example, CMV reactivation correlated with a myriad of proteins, including IL10, CXCL10, CXCL11, sCD40, sPDL1 (CD274), and sCD8A. Increases in sCD40, sPDL1, and sCD8A, in conjunction with the increased circulating activated CD4+ and CD8+ T-cells during CMV reactivation, suggests that CMV may elicit activation of T-cells, as previously reported^57^.

Similarly to CMV, EBV was also associated with a significant increase in plasma IL10, as has been observed in previous studies^58^. Increasing serum concentrations of IL10 may stem from both viral upregulation of human IL10, coupled with possible displacement of human IL10 from IL10R by virally produced IL10 competitive agonists encoded by both CMV and EBV^58–60^. Other than IL10, EBV was also associated with increases in four additional cytokines and chemokines: IL6, CXCL10, CCL7, and CCL2. IL6 and CXCL10 have been strongly correlated with COVID-19 severity ^34,36,61,62^, and given that EBV reactivation occurs early during acute COVID-19, these findings raise the possibility that EBV may play a role in the production of these cytokines in critically ill COVID-19 patients. Collectively, these findings suggest that viruses may either exploit existing excessive inflammation to reactivate, or magnify production of specific inflammatory cytokines, with both mechanisms potentially contributing to acute COVID-19 severity.

In addition to perturbations of cytokines, we also observed significant changes in the host transcriptome, particularly PBMCs, possibly reflecting responses to viral reactivation. Broadly, reactivation of *Herpesviridae* in the blood was associated with increased expression of genes involved in cellular replication, potentially mirroring the expanding lymphocyte response to the reactivation. In the nasal compartment, reactivation of *Herpesviridae* correlated with upregulation of localized inflammatory signaling.

We also observed associations between virome reactivation and levels of metabolites. In particular, levels of methylcysteine sulfoxide and 6-bromotryptophan, which are involved in protection against oxidative stress^63,64^, were negatively associated with detection of *Herpesviridae* transcripts, suggesting that viral reactivation may contribute to oxidative stress and cellular damage. Furthermore, 6-bromotryptophan has previously been associated with COVID-19 complications^34,65^ and impaired kidney function^63,66^, the persistently low levels of this metabolite in the context of viral reactivation could imply additional risk to kidney health, compounded by the setting of hospitalization. This finding aligns with the pronounced and sustained elevation of urea and TMAP in CMV-infected patients which is indicative of renal impairment and suggests the need for monitoring of CMV reactivation in COVID-19 patients presenting with acute kidney damage. Additionally, we observed elevations in long chain fatty acids and dimethylarginine with CMV reactivation, which have been previously implicated in higher inflammatory states during acute COVID-19^41,43^ and suggest a potential disturbance in endothelial function and systemic inflammation^42,44,67^. Collectively, these data suggest that viral reactivation is associated with specific metabolic perturbations, possibly reflecting strategies by these pathogens to manipulate host cell metabolism for their advantage^38^.

PASC, also known as Long COVID, is a disorder that is characterized by heterogeneous symptoms that persist for months to years’ post-acute COVID-19 infection, which can profoundly impact patients’ health and often results in disability and loss of income^45,68–70^. While true prevalence is unknown due to evolving definition of PASC and variability of study design across studies, it is estimated to be ∼10% or higher in patients post COVID-19^71–73^. Given the high prevalence and lack of treatments for PASC, there is an urgent need for a better understanding of the underlying PASC pathophysiology to guide the development of novel therapeutics. One of the leading emerging factors associated with PASC is chronic viral reactivation, particularly reactivation of EBV, for which PASC patients with neurocognitive symptoms have elevated antibody titers^22^.

In this study, we demonstrate that EBV reactivation is extensive in severe acute COVID-19 and establish the timing of EBV reactivation relative to SARS-CoV-2 infection and hospitalization. Although we did not find a direct association of EBV transcripts in the *acute* phase of COVID-19 with convalescent deficits, we observed a trend towards higher rates of PASC in participants with EBV reactivation and a trend toward lower rates in those with CMV reactivation, consistent with prior published reports which highlight the potential role of EBV reactivation in the development of PASC^22,26^. Of note, the PROs in this study were designed early in the pandemic prior to emergence of PASC, and thus may not fully capture PASC phenotypes, which may have limited our ability to detect association between EBV viral reactivation and PASC in our dataset. Furthermore, prior reports connecting EBV with PASC have focused strictly on EBV antibody titers, which was collected only at hospital admission for half our cohort (n=479). Thus, future work delving into the dynamics of antibody titers compared to viral transcription and reactivation may continue to elucidate EBV’s role in PASC.

Nonetheless, we report a novel significant association between the PASC Physical PRO group and detection of *Anelloviridae* transcripts in the convalescent period. *Anelloviridae* are a large family of negative-sense DNA stranded viruses, with some members of this family (e.g. Torque teno virus) found in ∼80-90% of the population^74,75^. Interestingly, *Anelloviridae* have been previously linked with chronic conditions, such as chronic fatigue syndrome and multiple sclerosis^76–78^. These conditions often present with physical symptoms similar to those reported by PASC patients, including fatigue, cognitive dysfunction, and post- exertional malaise. Thus, our findings suggest that detection of *Anelloviridae* transcripts may serve as a biomarker of persistent physical symptoms among PASC patients. Our data highlights the need for future research to dissect the role of viral reactivation, particularly of *Anelloviridae*, in the development and persistence of PASC, as well as to explore potential therapeutic interventions targeting this virus family in PASC patients.

Strengths of our study include a large multicenter prospective cohort, diverse bioassays enabling comprehensive immunophenotyping, detailed clinical phenotyping, and the use of RNA sequencing to measure actively replicating viruses. However, there were also several limitations to our study. First, the usage of transcripts to identify viral loads is seldom performed when compared to common clinical RT-qPCR tests. However, the extensive sequencing depth of our samples (targeted read depth 50,000,000 reads) provided sufficient depth to identify viral reads. Additionally, we had only six collection timepoints during the acute stage of disease, limiting the ability to absolutely rule out that participants did not have viral reactivation that might have occurred between collection timepoints. However, using the samples from >1000 participants and the fact that the exact date of samples varied from participant to participant due to the nature of observation and voluntary human studies, we calculated the overall global trends of reactivation over time. Another limitation was substantial participant drop out during the convalescent stage of the study, particularly in patients who had acute viral reactivation, particularly with EBV, limiting the power of analyses testing the association of acute viral reactivation with PASC. Lastly, IMPACC patients were unvaccinated and primarily exposed to ancestral strains of SARS-CoV-2, and thus further studies are needed to ascertain effects of more recent SARS-CoV-2 strains on viral reactivation in acute COVID-19 and PASC in population with hybrid immunity.

## Conclusion

In this study, we integrated clinical, immunologic, virologic, and multi-omic data from cellular and cytokine immunophenotyping, metabolomics, proteomics, and transcriptomics in a longitudinal cohort of >1000 COVID-19 patients to investigate for reactivation of chronic viral infections and their association with clinical outcomes in one of the largest studies to date. We found that multiple chronic viruses reactivate during acute COVID-19 infection, particularly from the *Herpesviridae* and *Anelloviridae* families. Furthermore, we delineate the temporal dynamics of reactivation for various viruses and report their associations with the host immune response, molecular pathways, as well as acute and chronic clinical sequelae of COVID-19. Notably, our results raise the possibility that viral reactivation may contribute to the development of PASC. This finding underscores the pressing need to address chronic viral reactivation in the evaluation and management of acute COVID-19 and PASC.

## Methods

### Study Design and Participant Recruitment

IMPACC is a prospective longitudinal study that enrolled >1,000 hospitalized COVID-19 patients, as previously described^34,45,62,79–81^. Participants 18 years and older were recruited from 20 hospitals across 15 academic institutes within the United States (Figure 1A). All participants were confirmed to be SARS-CoV-2 positive by reverse transcription PCR (RT-PCR) testing and no participants were vaccinated for SARS-CoV-2 at time of enrollment. Nasal swabs, blood, and endotracheal aspirate (for ventilated patients) were collected within 72 hours of hospital admission (visit 1) and on days 4, 7, 14, 21, and 28 post hospital admission in addition to convalescent samples at 3, 6, 9, and 12 months. Participants were characterized into one of five trajectory groups (TGs) based on latent mixed class modeling of a 7-point ordinal scale that characterized degree of respiratory illness and reflected acute COVID-19 severity^31^.

The Department of Health and Human Services Office for Human Research Protections (OHRP) and NIAID concurred that the IMPACC study qualified for public health surveillance exemption. The study protocol was sent for review to each site’s institutional review board (IRB), with twelve sites conducting as a public health surveillance study, and three sites integrating the IMPACC study into IRB-approved protocols (The University of Texas at Austin, IRB 2020-04- 0117; University of California San Francisco, IRB 20-30497; Case Western Reserve University, IRB STUDY20200573) with participants providing informed consent. Participants enrolled at sites operating as a public health surveillance study were provided information sheets describing the study including the samples to be collected and plans for analysis and data de-identification. Participants who requested not to participate after review of the study plan and information were not enrolled. Participants were not compensated while hospitalized but were subsequently compensated for outpatient visits and surveys. This study was registered at clinicaltrials.gov (NCT0438777) and followed the Strengthening the Reporting of Observational Studies in Epidemiology (STROBE) guidelines.

### Sample Processing and Assays

Samples were processed as previously described^34,45,62,79–81^, with the sample protocol extensively documented in the IMPACC study design and protocol paper^79^. Briefly, 10 mL of blood and nasal swabs were collected at each visit, with blood processed within 6 hours of collection. Blood was collected in both a 2.5 mL Greiner Vacuette CAT Serum Separating Tube (SST) (Cat: 454243P) for serum and a 7.5 mL Sarstedt Venous blood collection monovette EDTA (Cat: NC9453456) for whole blood, PBMCs, and plasma. The SST was kept vertical at room temperature (RT) for at least 30 minutes before centrifuging at RT for 10 minutes at 1000g. Serum was then aliquoted at 100 μL for downstream assays.

From the EDTA tube, it was briefly inverted to mix before aliquoting 270 uL of whole blood twice for both Cytometry Time of Flight (CyTOF) and genome wide association sequencing (GWAS) which was stored at –80C until shipment to their respective processing core. The remaining blood was centrifuged at RT for 10 minutes at 1000g before aliquoting and storing 500 uL of plasma at –80C for proteomic and metabolomics. PBMCs were then isolated from the remaining sample using the SepMate and Lymphoprep system (StemCell) following manufacturer protocol and as previously described^79^. PBMCs were then stored at 2.5 x 10^5^ cells in 200 μL of RLT Buffer (Qiagen) and beta-mercaptoethanol at –80C.

Interior nasal turbinate swabs (herein referred to as nasal swabs) were collected and stored in 1 mL of Zymo-DNA/RNA shield reagent (Zymo Research), before RNA was extracted twice in parallel from 250 μL of sample and purified with the KingFisher Flex sample purification system (ThermoFisher) and the quick DNA-RNA MagBead kit (Zymo Research). The duplicated RNA was pooled and aliquoted at 20 μL for the downstream assays (SARS-CoV-2 RT-qPCR and RNA-sequencing).

When participants were ventilated, an endotracheal aspirate (EA) was also collected in a 40 cc Argyle specimen trap and was processed within 2 hours of collection. First, 500 uL of 1:1 diluted EA with Maxpar PBS (Ca+2 -Mg+2 free) was mixed with 500 uL of DNA/RNA shield in a Zymo tube with lysis beads and subsequently stored at –80C for bulk RNA-sequencing.

Collected samples were then shipped and processed for nasal, PBMC, and EA RNA-sequencing, plasma proteomics, serum cytokine proximity extension assay (PEA), serum EBV and CMV antibody titers, whole blood CyTOF, and plasma metabolomics at their respective processing cores as previously described^34,45,62,79–81^. Each assay is described briefly below with additional technical details in the prior publications^34,45,62,79–81^.

- PBMC, EA, and Nasal RNA-seq were sequenced on a NovaSeq 6000 (Illumina) at 100 bp paired-end read length, and data was aligned using STAR (v2.4.2a or v2.4.3) against the GRCh38 reference genome. Gene counts were generated using HTSeq-count (v0.4.1).
- Nasal SARS-CoV-2 viral load was also measured by nasal swab RT-qPCR conducted using two separate sets of primers and probes for the N1 and N2 genes.
- Whole Blood CyTOF used a panel of 43 antibodies to quantify the frequency of 65 cell subsets using a Fluidigm Helios mass cytometer, with a semi-automatic gating strategy used for cell type assignment.
- Serum anti-viral antibodies for EBV and CMV were measured via a Luminex platform and were normalized to Assay Chex control beads by Radix BioSolutions with batch regressed as previously described^80^. This assay was only performed for half of the cohort (n=479).
- Plasma proteomics was evaluated with a EVOSEP one liquid chromatography connected to a TIMSTOF Pro (Bruker) as previously described^32,33^.
- Plasma metabolomics were measured using liquid chromatography-mass spectrometry and was conducted by Metabolon using in-house standards^34,82,83^.
- Serum cytokines and chemokines were measured using O-Link’s multiplex PEA for 92 proteins known to be involved in human inflammation (Olink Bioscience, Uppsala, Sweden).

Of note and unique to this IMPACC manuscript, taxonomic alignments for human infecting viruses from the PBMC, Nasal, and EA RNA-seq data was obtained from CZID^84^, which removes host reads before aligning remaining reads against the National Center for Biotechnology Information (NCBI) nucleotide and non-redundant databases. A sample was considered “positive” for a virus if it had at least one read that mapped to both the nucleotide and non-redundant database. No water control samples from any of the RNA-sequencing had any reads for the human-infecting viruses evaluated in this manuscript, supporting this low threshold for positivity.

### Statistics

All analyses were executed in R v4.0.3. All p-values calculated in this manuscript were adjusted using the Benjamini-Hochberg procedure where appropriate and are indicated by “adj.p” or “adjusted p-value” (circumstances where no adjustments were necessary will instead report “p” or “p-value”).

#### Clinical Features and Demographics

For testing the association of detection of chronic viral transcripts with TGs and age, we used cumulative linked mixed modeling from the ordinal (v 2019.12-10) R package. Due to both COVID-19 severity and age quantiles being ordinal, cumulative linked mixed modeling allowed for this ordinal relationship to be accounted for in addition to including enrollment site as a random effect. In the age quantile ordinal model, trajectory groups were also used as a main effect to control for COVID-19 severity.

Trajectory_group ∼ virus_status, random = enrollment_site

Admit_age_quintile ∼ virus_status + trajectory_group, random = enrollment_site

For association testing of specific clinical complications, comorbidities, medication, and other clinical outcomes, linear mixed effect modeling from the lme4 R package (v1.1-28) was used with viral status, sex, admit_age_quintile, and trajectory group as main effects and enrollment site as a random effect.

Clinical_feature ∼ virus_status + sex + admit_age_quintile + trajectory_group + (1 | enrollment_site)

#### Serum Cytokines and Plasma Metabolomics

To evaluate both the serum PEA cytokine assay and the plasma metabolomics, generalized additive mixed modeling (gamm) from the gamm4 (v 0.2-6) R package was used to evaluate for differences in individual analytes between patients who had detected transcripts for a chronic virus compared to patients who never had human-infecting viral transcripts detected (other than for SARS-CoV-2). Analytes were modeled against days from admission using cubic regression splines with interactions of both status for a given chronic virus (binary: positive or negative based on detected transcripts at any collected timepoint) and TG in addition to fixed effects of status for the chronic virus being evaluated, TG, sex, age at time of admission sorted into quintiles, and SARS-CoV-2 nasal viral RPM at the time of that sample. As chronic viral status is both a main effect and interaction term in the model, we used a lower adjusted p-value cutoff of 0.01 to account for the fact that a feature was significant if either term was significant.

Analyte ∼ s(days, bs = ‘cr’) + s(days, bs = ‘cr’, by = ‘virus_status’) + s(days, bs = ‘cr’, by = ‘trajectory_group’) + virus_status + sex + trajectory_group + admit_age_quintile + sarscovs2_nasal_log_rpm, random = ∼(1|enrollment_site/participant_id)

#### Nasal and PBMC RNA-Sequencing

To analyze the signature of host gene expression associated with chronic viruses, the limma (v 3.46.0) R package was used for both the nasal and PBMC RNA-sequencing to evaluate differential expressed genes associated with participants who had detectable chronic viral transcripts.

∼ admit_age_quintile + sex + visit_number + trajectory_group + sarscov2_nasal_log_rpm + virus_status

Pathway enrichment on both the upregulated and downregulated differentially expressed genes was conducted using hypergeometric enrichment testing from the R package clusterProfiler (v3.18.1) with the Reactome pathway database.

### Data and code availability

Data used in this study is available at ImmPort Shared Data under the accession number <u>SDY1760</u> and in the NLM’s Database of Genotypes and Phenotypes (dbGaP) under the accession number <u>phs002686.v2.p2</u>. All code is deposited on Bitbucket (https://bitbucket.org/kleinstein/impacc-public-code/src/master/chronic_viruses_manuscript/).

## IMPACC Network Competing Interests

The Icahn School of Medicine at Mount Sinai has filed patent applications relating to SARS-CoV-2 serological assays and NDV-based SARS-CoV-2 vaccines which list Florian Krammer as co-inventor. Mount Sinai has spun out a company, Kantaro, to market serological tests for SARS-CoV-2. Florian Krammer has consulted for Merck and Pfizer (before 2020), and is currently consulting for Pfizer, Seqirus, 3rd Rock Ventures, Merck and Avimex. The Krammer laboratory is also collaborating with Pfizer on animal models of SARS-CoV-2. Viviana Simon is a co-inventor on a patent filed relating to SARS-CoV-2 serological assays (the “Serology Assays”). Ofer Levy is a named inventor on patents held by Boston Children’s Hospital relating to vaccine adjuvants and human in vitro platforms that model vaccine action. His laboratory has received research support from GlaxoSmithKline (GSK) and is a co-founder of and advisor to Ovax, Inc. Charles Cairns serves as a consultant to bioMerieux and is funded for a grant from Bill & Melinda Gates Foundation. James A Overton is a consultant at Knocean Inc. Jessica Lasky-Su serves as a scientific advisor of Precion Inc. Scott R. Hutton, Greg Michelloti and Kari Wong are employees of Metabolon Inc. Vicki Seyfert-Margolis is a current employee of MyOwnMed. Nadine Rouphael reports grants or contracts with Merck, Sanofi, Pfizer, Vaccine Company, Quidel, Lilly and Immorna, and has participated on data safety monitoring boards for Moderna, Sanofi, Seqirus, Pfizer, EMMES, ICON, BARDA, Imunon, CyanVac and Micron. Nadine Rouphael has also received support for meetings/travel from Sanofi and Moderna and honoraria from Virology Education. Adeeb Rahman is a current employee of Immunai Inc. Steven Kleinstein is a consultant related to ImmPort data repository for Peraton. Nathan Grabaugh is a consultant for Tempus Labs and the National Basketball Association. Akiko Iwasaki is a consultant for 4BIO, Blue Willow Biologics, Revelar Biotherapeutics, RIGImmune, Xanadu Bio, Paratus Sciences. Monika Kraft receives research funds paid to her institution from NIH, ALA; Sanofi, Astra-Zeneca for work in asthma, serves as a consultant for Astra-Zeneca, Sanofi, Chiesi, GSK for severe asthma; is a co-founder and CMO for RaeSedo, Inc, a company created to develop peptidomimetics for treatment of inflammatory lung disease. Esther Melamed received research funding from Babson Diagnostics and honorarium from Multiple Sclerosis Association of America and has served on the advisory boards of Genentech, Horizon, Teva, and Viela Bio. Carolyn Calfee receives research funding from NIH, FDA, DOD, Roche-Genentech and Quantum Leap Healthcare Collaborative as well as consulting services for Janssen, Vasomune, Gen1e Life Sciences, NGMBio, and Cellenkos. Wade Schulz was an investigator for a research agreement, through Yale University, from the Shenzhen Center for Health Information for work to advance intelligent disease prevention and health promotion; collaborates with the National Center for Cardiovascular Diseases in Beijing; is a technical consultant to Hugo Health, a personal health information platform; cofounder of Refactor Health, an AI-augmented data management platform for health care; and has received grants from Merck and Regeneron Pharmaceutical for research related to COVID-19. Grace A McComsey received research grants from Redhill, Cognivue, Pfizer, and Genentech, and served as a research consultant for Gilead, Merck, Viiv/GSK, and Jenssen. Linda N. Geng received research funding paid to her institution from Pfizer, Inc.

## Author Contributions

CM, JC, NR, HCP, VP, KH, AG, VC, RD, the IMPACC Network, KKS, EFR, OL, HM, PH, HS, JDA, CRL, and EM conceived the study. CM, JC, and KH conducted formal analysis. CM and JC designed software. CM, CRL, and EM designed the study methodology. The IMPACC Network acquired funding. Supervision was provided by NR, KKS, EFR, OL, HM, PH, HS, JDA, CRL, and EM. All authors edited and reviewed the manuscript.

## Acknowledgment

We thank the participants of the study for their voluntary enrollment and contribution of samples for this work. See the supplement for details on the IMPACC Network. We acknowledge the assistance of the following individuals: Sanya Thomas, Mitchell Cooney, Shun Rao, Sofia Vignolo, and Elena Morrocchi (all from the CDCC); Arash Naeim, Marianne Bernardo, Sarahmay Sanchez, Shannon Intluxay, Clara Magyar, Jenny Brook, Estefania Ramires-Sanchez, Megan Llamas, Claudia Perdomo, Clara E. Magyar, and Jennifer A. Fulcher (all from the David Geffen School of Medicine at UCLA); members of the UCLA Center for Pathology Research Services and the Pathology Research Portal; M. Catherine Muenker, Dimitri Duvilaire, Maxine Kuang, William Ruff, Khadir Raddassi, Denise Shepherd, Haowei Wang, Omkar Chaudhary, Syim Salahuddin, John Fournier, Michael Rainone, and Maxine Kuang (all from the Yale School of Medicine). We thank the leadership of Boston Children’s Hospital including Drs. Wendy Chung, Gary Fleisher and Kevin Churchwell for their support for the Precision Vaccines Program. Dr. Augustine’s and Becker’s co-authorship of this report does not necessarily represent the official views of the National Institute of Allergy and Infectious Diseases, the National Institutes of Health or any other agency of the United States Government.

\# The IMPACC Network

**National Institute of Allergy and Infectious Diseases, National Institute of Health, Bethesda, MD 20814, USA:** Patrice M. Becker, Alison D. Augustine, Steven M. Holland, Lindsey B. Rosen, Serena Lee, Tatyana Vaysman

**Clinical and Data Coordinating Center (CDCC) Precision Vaccines Program, Boston Children’s Hospital, Boston, MA 02115, USA:** Al Ozonoff, Joann Diray-Arce, Jing Chen, Alvin Kho, Carly E. Milliren, Annmarie Hoch, Ana C. Chang, Kerry McEnaney, Brenda Barton, Claudia Lentucci, Maimouna D. Murphy, Mehmet Saluvan, Tanzia Shaheen, Shanshan Liu, Caitlin Syphurs, Marisa Albert, Arash Nemati Hayati, Robert Bryant, James Abraham, Sanya Thomas, Mitchell Cooney, Meagan Karoly

Benaroya Research Institute, University of Washington, Seattle, WA 98101, USA: **Matthew**

1. C. Altman, Naresh Doni Jayavelu, Scott Presnell, Bernard Kohr, Tomasz Jancsyk, Azlann Arnett

**La Jolla Institute for Immunology, La Jolla, CA 92037, USA:** Bjoern Peters, James A. Overton, Randi Vita, Kerstin Westendorf

**Knocean Inc. Toronto, ON M6P 2T3, Canada:** James A. Overton

**Precision Vaccines Program, Boston Children’s Hospital, Harvard Medical School, Boston, MA 02115, USA:** Ofer Levy, Hanno Steen, Patrick van Zalm, Benoit Fatou, Kinga K. Smolen, Arthur Viode, Simon van Haren, Meenakshi Jha, David Stevenson, Oludare Odumade

**Brigham and Women’s Hospital, Harvard Medical School, Boston, MA 02115, USA:** Lindsey R. Baden, Kevin Mendez, Jessica Lasky-Su, Alexandra Tong, Rebecca Rooks, Michael Desjardins, Amy C. Sherman, Stephen R. Walsh, Xhoi Mitre, Jessica Cauley, Xiofang Li, Bethany Evans, Christina Montesano, Jose Humberto Licona, Jonathan Krauss, Nicholas C. Issa, Jun Bai Park Chang, Natalie Izaguirre

**Metabolon Inc, Morrisville, NC 27560, USA:** Scott R. Hutton, Greg Michelotti, Kari Wong

**Prevention of Organ Failure (PROOF) Centre of Excellence, University of British Columbia, Vancouver, BC V6T 1Z3, Canada:** Scott J. Tebbutt, Casey P. Shannon

**Case Western Reserve University and University Hospitals of Cleveland, Cleveland, OH 44106, USA:** Rafick-Pierre Sekaly, Slim Fourati, Grace A. McComsey, Paul Harris, Scott Sieg, Susan Pereira Ribeiro

**Drexel University, Tower Health Hospital, Philadelphia, PA 19104, USA:** Charles B. Cairns, Elias K. Haddad, Michele A. Kutzler, Mariana Bernui, Gina Cusimano, Jennifer Connors, Kyra Woloszczuk, David Joyner, Carolyn Edwards, Edward Lee, Edward Lin, Nataliya Melnyk, Debra

1. L. Powell, James N. Kim, I. Michael Goonewardene, Brent Simmons, Cecilia M. Smith, Mark Martens, Brett Croen, Nicholas C. Semenza, Mathew R. Bell, Sara Furukawa, Renee McLin, George P. Tegos, Brandon Rogowski, Nathan Mege, Kristen Ulring, Pam Schearer, Judie Sheidy, Crystal Nagle

**MyOwnMed Inc., Bethesda, MD 20817, USA:** Vicki Seyfert-Margolis

**Emory School of Medicine, Atlanta, GA 30322, USA:** Nadine Rouphael, Steven E. Bosinger, Arun K. Boddapati, Greg K. Tharp, Kathryn L. Pellegrini, Brandi Johnson, Bernadine

Panganiban, Christopher Huerta, Evan J. Anderson, Hady Samaha, Jonathan E. Sevransky, Laurel Bristow, Elizabeth Beagle, David Cowan, Sydney Hamilton, Thomas Hodder, Amer Bechnak, Andrew Cheng, Aneesh Mehta, Caroline R. Ciric, Christine Spainhour, Erin Carter, Erin M. Scherer, Jacob Usher, Kieffer Hellmeister, Laila Hussaini, Lauren Hewitt, Nina Mcnair, Susan Pereira Ribeiro, Sonia Wimalasena

**Icahn School of Medicine at Mount Sinai, New York, NY 10029, USA:** Ana Fernandez-Sesma, Viviana Simon, Florian Krammer, Harm Van Bakel, Seunghee Kim-Schulze, Ana Silvia Gonzalez Reiche, Jingjing Qi, Brian Lee, Juan Manuel Carreño, Gagandeep Singh, Ariel Raskin, Johnstone Tcheou, Zain Khalil, Adriana van de Guchte, Keith Farrugia, Zenab Khan, Geoffrey Kelly, Komal Srivastava, Lily Q. Eaker, Maria C. Bermúdez-González, Lubbertus C.F. Mulder, Katherine F. Beach, Miti Saksena, Deena Altman, Erna Kojic, Levy A. Sominsky, Arman Azad, Dominika Bielak, Hisaaki Kawabata, Temima Yellin, Miriam Fried, Leeba Sullivan, Sara Morris, Giulio Kleiner, Daniel Stadlbauer, Jayeeta Dutta, Hui Xie, Manishkumar Patel, Kai Nie

**Immunai Inc. New York, NY 10016, USA:** Adeeb Rahman **Oregon Health Sciences University, Portland, OR 97239, USA:** William B. Messer, Catherine 1. L. Hough, Sarah A.R. Siegel, Peter E. Sullivan, Zhengchun Lu, Amanda E. Brunton, Matthew Strand, Zoe L. Lyski, Felicity J. Coulter, Courtney Micheleti

**Stanford University School of Medicine, Palo Alto, CA 94305, USA:** Holden Maecker, Bali Pulendran, Kari C. Nadeau, Yael Rosenberg-Hasson, Michael Leipold, Natalia Sigal, Angela Rogers, Andrea Fernandes, Monali Manohar, Evan Do, Iris Chang, Alexandra S. Lee, Catherine Blish, Henna Naz Din, Jonasel Roque, Linda Geng, Maja Artandi, Mark M. Davis, Neera Ahuja, Samuel S. Yang, Sharon Chinthrajah, Thomas Hagan

**David Geffen School of Medicine at the University of California Los Angeles, Los Angeles CA 90095, USA:** Elaine F. Reed, Joanna Schaenman, Ramin Salehi-Rad, Adreanne M. Rivera, Harry C. Pickering, Subha Sen, David Elashoff, Dawn C. Ward, Jenny Brook, Estefania Ramires Sanchez, Megan Llamas, Claudia Perdomo, Clara E. Magyar, Jennifer Fulcher

**University of California San Francisco, San Francisco, CA 94115, USA:** David J. Erle, Carolyn S. Calfee, Carolyn M. Hendrickson, Kirsten N. Kangelaris, Viet Nguyen, Deanna Lee, Suzanna Chak, Rajani Ghale, Ana Gonzalez, Alejandra Jauregui, Carolyn Leroux, Luz Torres Altamirano, Ahmad Sadeed Rashid, Andrew Willmore, Prescott G. Woodruff, Matthew F. Krummel, Sidney Carrillo, Alyssa Ward, Charles R. Langelier, Ravi Patel, Michael Wilson, Ravi Dandekar, Bonny Alvarenga, Jayant Rajan, Walter Eckalbar, Andrew W. Schroeder, Gabriela K. Fragiadakis, Alexandra Tsitsiklis, Eran Mick, Yanedth Sanchez Guerrero, Christina Love, Lenka Maliskova, Michael Adkisson, Aleksandra Leligdowicz, Alexander Beagle, Arjun Rao, Austin Sigman, Bushra Samad, Cindy Curiel, Cole Shaw, Gayelan Tietje-Ulrich, Jeff Milush, Jonathan Singer, Joshua J. Vasquez, Kevin Tang, Legna Betancourt, Lekshmi Santhosh, Logan Pierce, Maria Tecero Paz, Michael Matthay, Neeta Thakur, Nicklaus Rodriguez, Nicole Sutter, Norman Jones, Pratik Sinha, Priya Prasad, Raphael Lota, Saurabh Asthana, Sharvari Bhide, Tasha Lea, Yumiko Abe-Jones

**Yale School of Medicine, New Haven, CT 06510, USA:** David A. Hafler, Ruth R. Montgomery, Albert C. Shaw, Steven H. Kleinstein, Jeremy P. Gygi, Shrikant Pawar, Anna Konstorum, Ernie Chen, Chris Cotsapas, Xiaomei Wang, Leqi Xu, Charles Dela Cruz, Akiko Iwasaki, Subhasis Mohanty, Allison Nelson, Yujiao Zhao, Shelli Farhadian, Hiromitsu Asashima, Omkar Chaudhary,

Andreas Coppi, John Fournier, M. Catherine Muenker, Allison Nelson, Khadir Raddassi, Michael Rainone, William Ruff, Syim Salahuddin, Wade L. Shulz, Pavithra Vijayakumar, Haowei Wang, Esio Wunder Jr., H. Patrick Young, Albert I. Ko,

**Yale School of Public Health, New Haven, CT 06510, USA:** Denise Esserman, Leying Guan, Anderson Brito, Jessica Rothman, Nathan D. Grubaugh

**Baylor College of Medicine and the Center for Translational Research on Inflammatory Diseases, Houston, TX 77030, USA:** David B. Corry, Farrah Kheradmand, Li-Zhen Song, Ebony Nelson

**Oklahoma University Health Sciences Center, Oklahoma City, OK 73104, USA:** Jordan P. Metcalf, Nelson I. Agudelo Higuita, Lauren A. Sinko, J. Leland Booth, Douglas A. Drevets, Brent 1. R. Brown

**University of Arizona, Tucson AZ 85721, USA:** Monica Kraft, Chris Bime, Jarrod Mosier, Heidi Erickson, Ron Schunk, Hiroki Kimura, Michelle Conway, Dave Francisco, Allyson Molzahn, Connie Cathleen Wilson, Ron Schunk, Trina Hughes, Bianca Sierra

**University of Florida, Gainesville, FL 32611, USA:** Mark A. Atkinson, Scott C. Brakenridge, Ricardo F. Ungaro, Brittany Roth Manning, Lyle Moldawer University of Florida, Jacksonville, FL 32218, USA: Jordan Oberhaus, Faheem W. Guirgis University of South Florida, Tampa FL 33620, USA: Brittney Borresen, Matthew L. Anderson

**The University of Texas at Austin, Austin, TX 78712, USA:** Lauren I. R. Ehrlich, Esther Melamed, Cole Maguire, Dennis Wylie, Justin F. Rousseau, Kerin C. Hurley, Janelle N. Geltman, Nadia Siles, Jacob E. Rogers, Pablo Guaman Tipan.

## IMPACC Network Competing Interests

The Icahn School of Medicine at Mount Sinai has filed patent applications relating to SARS-CoV-2 serological assays and NDV-based SARS-CoV-2 vaccines which list Florian Krammer as co-inventor. Mount Sinai has spun out a company, Kantaro, to market serological tests for SARS-CoV-2. Florian Krammer has consulted for Merck and Pfizer (before 2020), and is currently consulting for Pfizer, Seqirus, 3rd Rock Ventures, Merck and Avimex. The Krammer laboratory is also collaborating with Pfizer on animal models of SARS-CoV-2. Viviana Simon is a co-inventor on a patent filed relating to SARS-CoV-2 serological assays (the “Serology Assays”). Ofer Levy is a named inventor on patents held by Boston Children’s Hospital relating to vaccine adjuvants and human in vitro platforms that model vaccine action. His laboratory has received research support from GlaxoSmithKline (GSK) and is a co-founder of and advisor to Ovax, Inc. Charles Cairns serves as a consultant to bioMerieux and is funded for a grant from Bill & Melinda Gates Foundation. James A Overton is a consultant at Knocean Inc. Jessica Lasky-Su serves as a scientific advisor of Precion Inc. Scott R. Hutton, Greg Michelloti and Kari Wong are employees of Metabolon Inc. Vicki Seyfert-Margolis is a current employee of MyOwnMed. Nadine Rouphael reports grants or contracts with Merck, Sanofi, Pfizer, Vaccine Company, Quidel, Lilly and Immorna, and has participated on data safety monitoring boards for Moderna, Sanofi, Seqirus, Pfizer, EMMES, ICON, BARDA, Imunon, CyanVac and Micron.

Nadine Rouphael has also received support for meetings/travel from Sanofi and Moderna and honoraria from Virology Education. Adeeb Rahman is a current employee of Immunai Inc.

Steven Kleinstein is a consultant related to ImmPort data repository for Peraton. Nathan

Grabaugh is a consultant for Tempus Labs and the National Basketball Association. Akiko Iwasaki is a consultant for 4BIO, Blue Willow Biologics, Revelar Biotherapeutics, RIGImmune, Xanadu Bio, Paratus Sciences. Monika Kraft receives research funds paid to her institution from NIH, ALA; Sanofi, Astra-Zeneca for work in asthma, serves as a consultant for Astra-Zeneca, Sanofi, Chiesi, GSK for severe asthma; is a co-founder and CMO for RaeSedo, Inc, a company created to develop peptidomimetics for treatment of inflammatory lung disease. Esther Melamed received research funding from Babson Diagnostics and honorarium from Multiple Sclerosis Association of America and has served on the advisory boards of Genentech, Horizon, Teva, and Viela Bio. Carolyn Calfee receives research funding from NIH, FDA, DOD, Roche-Genentech and Quantum Leap Healthcare Collaborative as well as consulting services for Janssen, Vasomune, Gen1e Life Sciences, NGMBio, and Cellenkos. Wade Schulz was an investigator for a research agreement, through Yale University, from the Shenzhen Center for Health Information for work to advance intelligent disease prevention and health promotion; collaborates with the National Center for Cardiovascular Diseases in Beijing; is a technical consultant to Hugo Health, a personal health information platform; cofounder of Refactor Health, an AI-augmented data management platform for health care; and has received grants from Merck and Regeneron Pharmaceutical for research related to COVID-19. Grace A McComsey received research grants from Redhill, Cognivue, Pfizer, and Genentech, and served as a research consultant for Gilead, Merck, Viiv/GSK, and Jenssen. Linda N. Geng received research funding paid to her institution from Pfizer, Inc.

