## Supplemental Figures for "Chronic Viral Reactivation and Associated Host Immune Response and Clinical Outcomes in Acute COVID-19 and Post-Acute Sequelae of COVID-19"

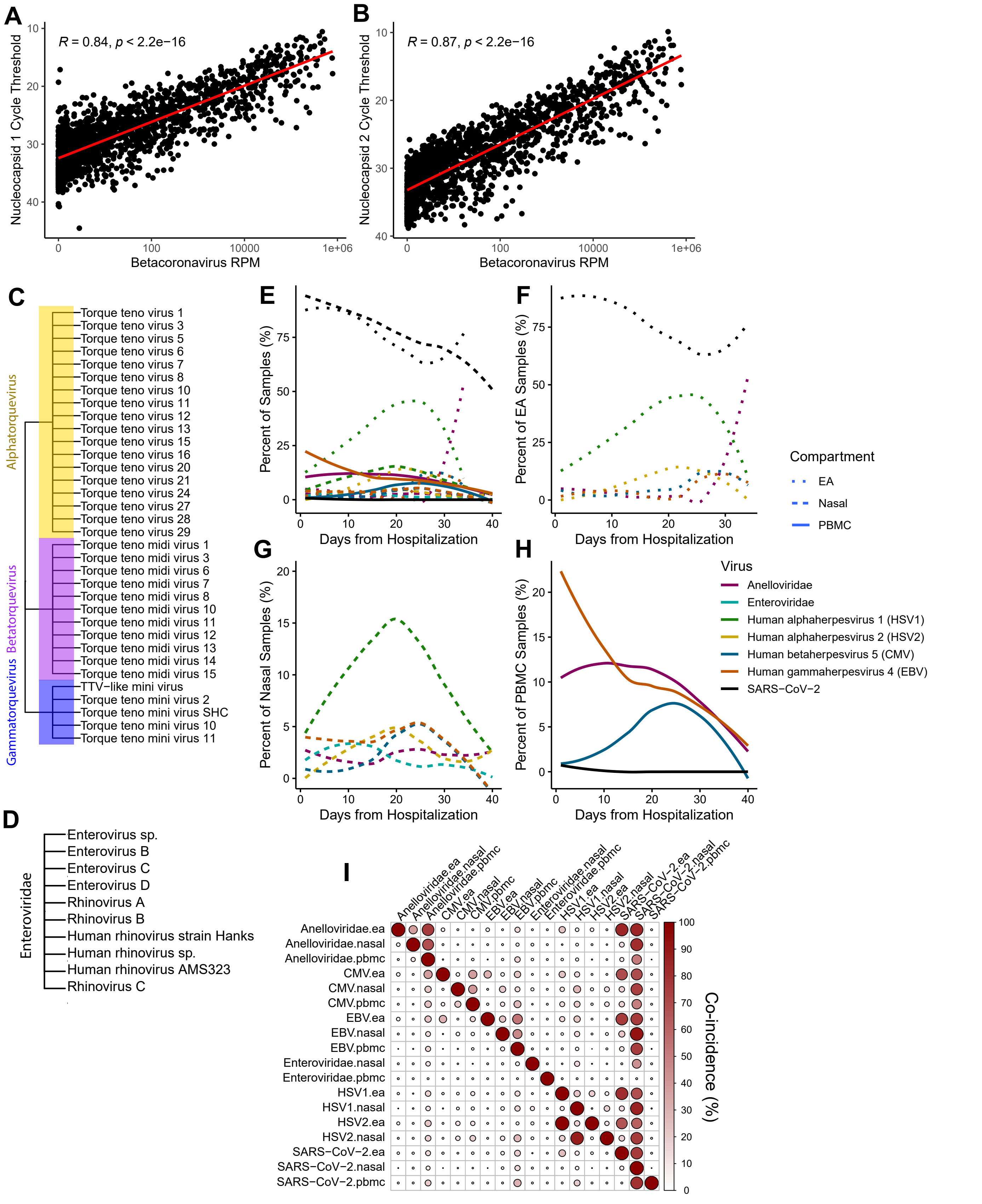


**Supplemental Figure 1 (related to Figure 1) – Diverse Chronic Viral Reactivation Occurs in Acute COVID-19.** Correlation of Betacoronavirus reads per million (RPM) from the nasal transcriptomics with **A)** Nucleocapsid 1 cycle threshold and **B)** Nucleocapsid 2 cycle threshold. **C)** Phylogenetic tree of the detected Anelloviridae in the PBMC transcriptomics (these were collectively collapsed in Anelloviridae for all analyses). **D)** Phylogenetic tree of the detected Enteroviruses in the nasal transcriptomics (these were collectively collapsed in Enterovirus for all analyses). **E)** Smoothed curves demonstrating the proportion of total samples that were positive for the five most common viruses in the nasal, PBMC, and EA transcriptomics. Curves were calculated by the proportion of samples positive for each day ± two days (rolling window), followed by a local polynomial regression fitting. **F)** (E) filtered to only the EA viruses. **G)** (E) filtered to only the nasal viruses. **H)** (E) filtered to only the PBMC viruses. **I)** A conditional probability table that depicts the percent of patient visits that the virus indicated on the column is found when the virus on the row is present.

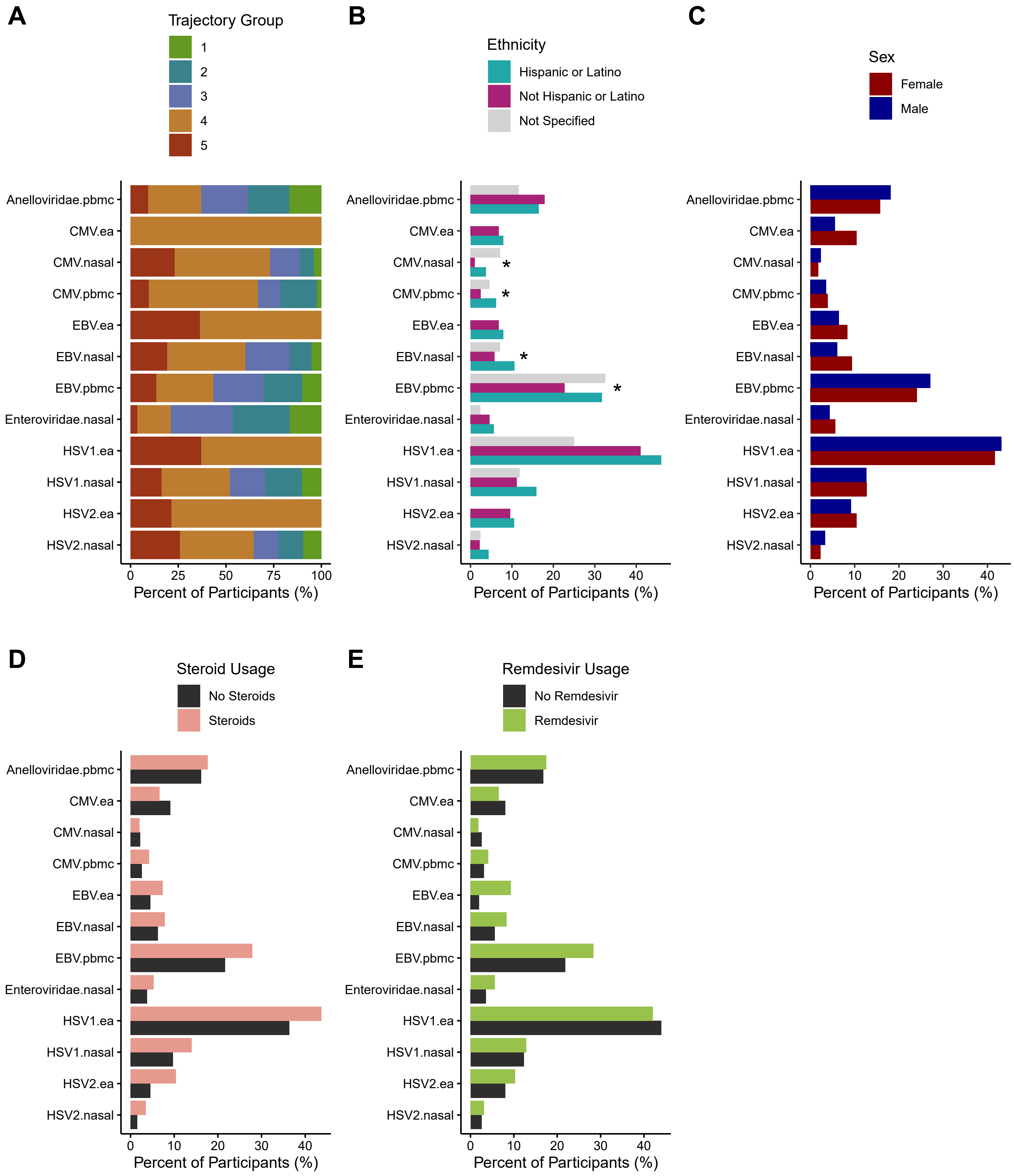


**Supplemental Figure 2 (related to Figure 2) – Additional Clinical Demographics and Medication Usage's Relationship with Chronic Viral Reactivation. A)** Percent of patients positive for each virus and compartment within the first 40 days post-hospitalization that are in each trajectory group (TG). Percent of participants positive for each virus in the respective compartments for any sample within the first 40 days post-hospitalization split by **B)** ethnicity, **C)** sex, **D)** administration of steroids during acute stay, **E)** administration of Remdesivir during acute COVID-19. Significance calculated using chi-square test of independence with p-value corrections following Benjamini-Hockberg procedure, * indicates adj.p<0.05.

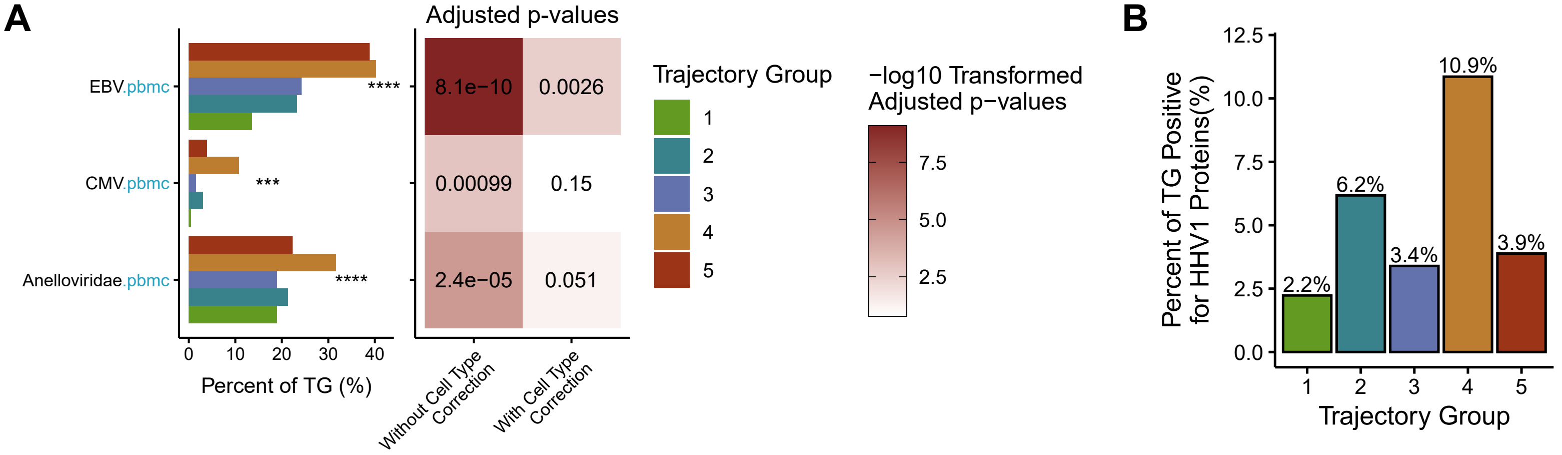


**Supplemental Figure 3 (related to Figure 3) – EBV Associates with COVID-19 Severity Even While Controlling for Changes in Circulating Blood Cell Composition. A)** Percent of participants in the cohort who had detectable viral reads in at least one sample within 40 days of hospital admission for the respective virus in the PBMC split by each trajectory group (on the left). Right: P-value for association of viral prevalence with TG both before and after correcting for circulating levels of significantly associated cell types (as determined from Figure 3A) from whole blood CyTOF. **B)** Percent of each TG with detectable HHV1 proteins in plasma via mass spectrometry at any sample within 40 days of hospital admission.

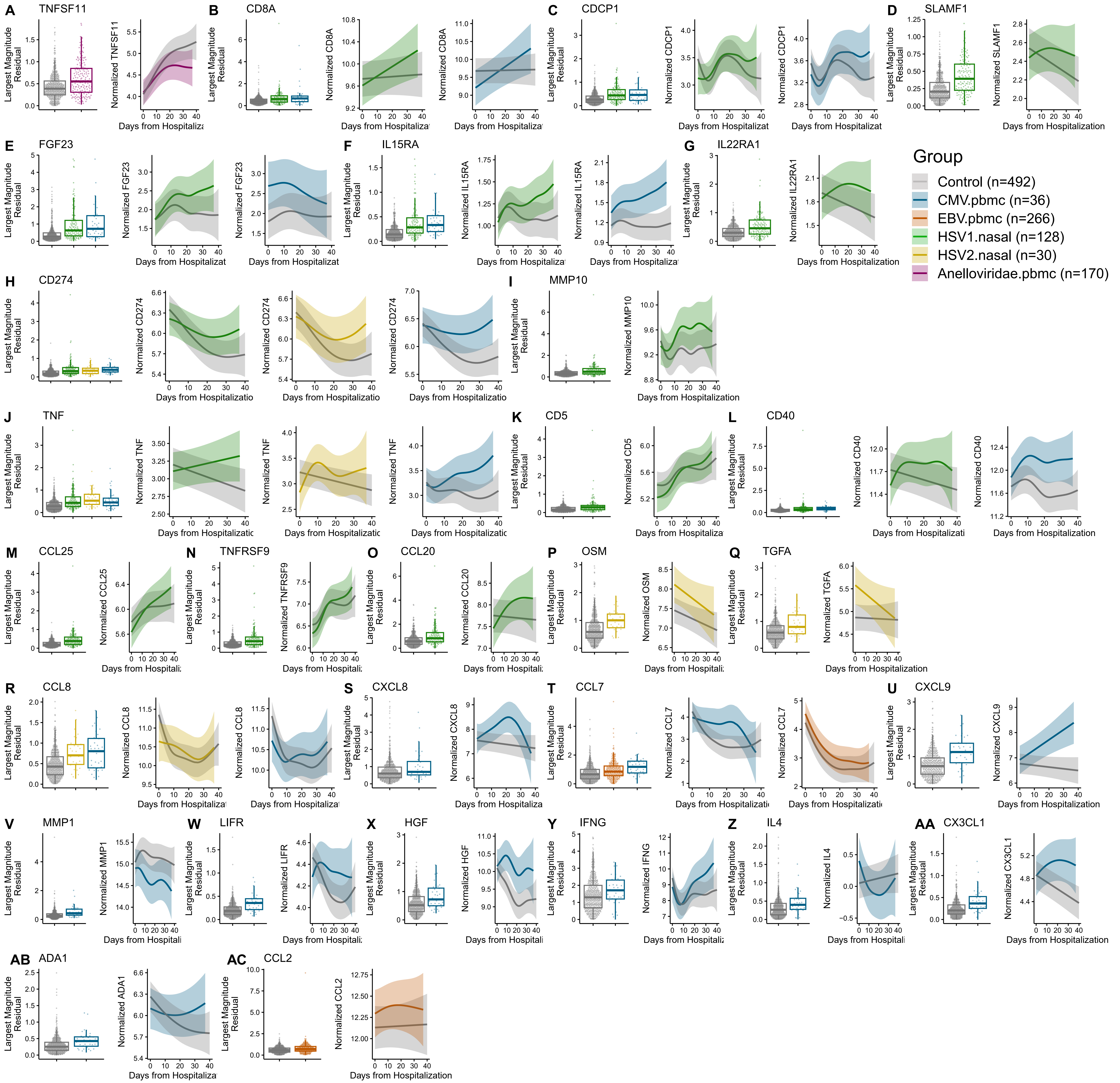


**Supplemental Figure 4 (related to Figure 4) – Serum Cytokines, Chemokines, and Soluble Inflammatory Proteins Trajectories in Acute COVID-19 Altered by Various Chronic Viral Reactivations.** Additional plots visualizing the largest magnitude gamm residual for each participant and longitudinal 95% confidence interval for cytokines not visualized in Figure 4.

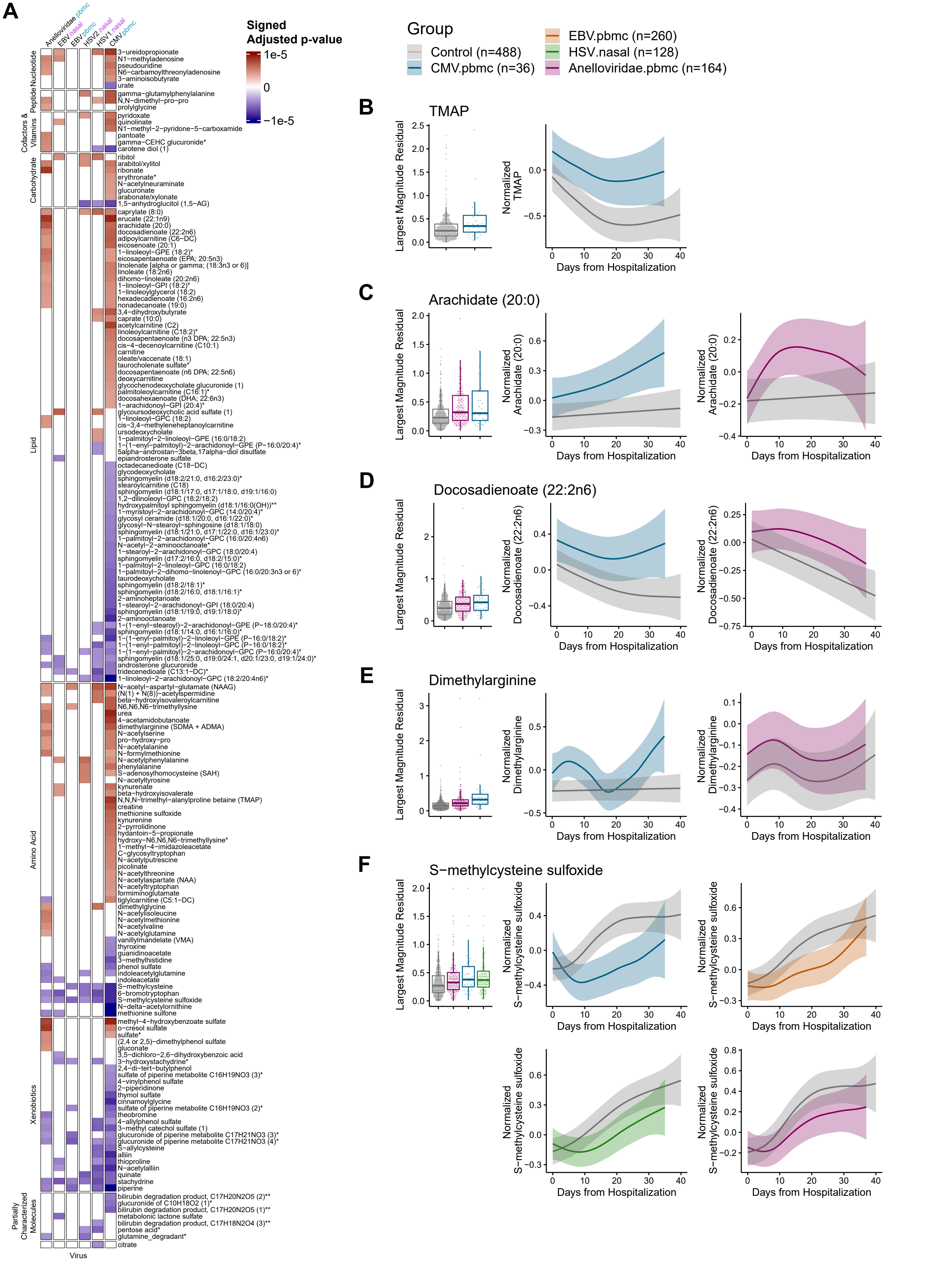


**Supplemental Figure 5 (related to Figure 5) – Chronic Viral Reactivation Correlates with Widespread Changes in the Metabolome. A)** Heatmap of all significantly associated metabolites for the different chronic viruses in their respective tissue compartments (i.e. Nasal or PBMC). Increasingly red represents a more significant positive association and increasingly blue represents an increasingly significant negative association. Largest magnitude gamm residual for each participant and longitudinal 95% confidence interval for **B)** TMAP, **C)** Arachidate, and **D)** Docosadianoate, **E)** Dimethylarginine, and **F**) S-methylcysteine sulfoxide.

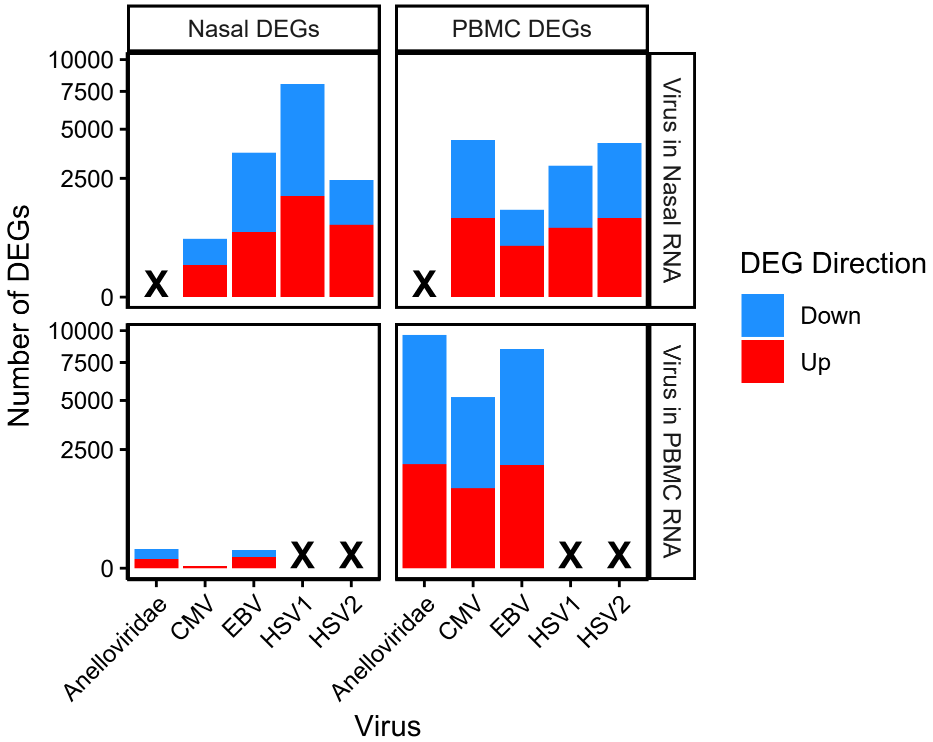


**Supplemental Figure 6 (related to Figure 6) – Location of Viral Reactivation Determines which Host Compartment will have Changes in Gene Expression.** Number of differentially expressed genes (DEGs) for non-SARS-CoV-2 viruses detected. The graph is split by the number of DEGs detected for the nasal host gene expression and PBMC host gene expression (columns) as well as by which tissue/sample the virus was detected (rows). Xs denote virus was not detected frequently enough in the tissue for analysis (i.e. HSV1 and HSV2 in the PBMCs and Anelloviridae in the Nasal). This plot demonstrates that viral reactivation in the nasal compartment affects both the Nasal and PBMC host gene expression, whereas reactivation in the PBMC only substantially affects gene expression in the PBMCs.

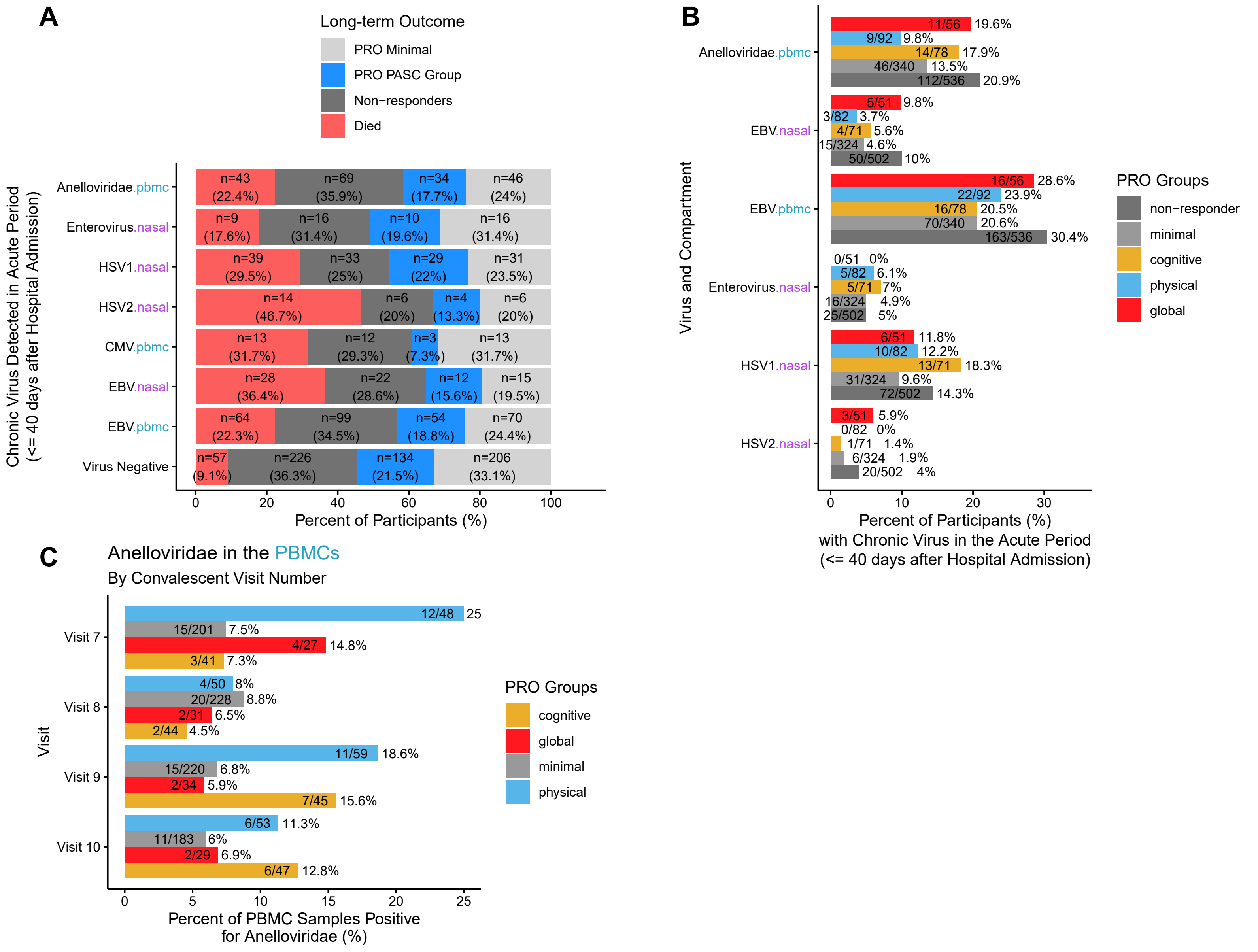


**Supplemental Figure 7 (related to Figure 7) – Chronic Viral Reactivation Correlates with Lower Responses on Convalescent Surveys and Anelloviridae are more Frequent in Convalescent Samples from the Physical PRO Group. A)** Percent of participants who either died, did not respond to convalescent symptom surveys (non-responders), or did respond and were grouped with a PASC affiliated PRO group or the PRO minimal group displaying minimal deficits. **B)** Percent of each PRO group and non-responders that had viruses detected in the acute COVID-19 periods. **C)** Percent of PRO group that had Anelloviridae detected in the outpatient convalescent samples, split by visit.
